# Neurophysiological Correlates of the Dunning-Kruger Effect

**DOI:** 10.1101/2019.12.26.888511

**Authors:** Alana Muller, Lindsey A. Sirianni, Richard J. Addante

**Author notes:** Corresponding Author: Richard J. Addante, PhD, California State University – San Bernardino Department of Psychology, 5500 University Parkway, San Bernardino, CA 92407, USA.

## Abstract

The Dunning-Kruger Effect (DKE) is a metacognitive phenomenon of illusory superiority in which individuals who perform poorly on a task believe they performed better than others, yet individuals who performed very well believe they under-performed compared to others. This phenomenon has yet to be directly explored in episodic memory, nor explored for reaction times or physiological correlates. We designed a novel method to elicit the DKE via a test of item recognition while electroencephalography (EEG) was recorded. Throughout the task, participants were asked to estimate the percentile in which they performed compared to others. Results revealed participants in the bottom 25th percentile overestimated their percentile, while participants in the top 75th percentile underestimated their percentile, exhibiting the classic DKE. Reaction time measures revealed a condition x group interaction whereby over-estimators responded faster than under-estimators when estimating being in the top percentile and responded slower when estimating being in the bottom percentile.

Between-group EEG differences were evident between over-estimators and under-estimators during Dunning-Kruger responses, which revealed FN400-like effects of familiarity supporting differences for over-estimators from 400-600 ms, whereas ‘old-new’ memory ERP effects revealed a late parietal component (LPC) associated with recollection-based processing from 600-900 ms for under-estimators that was not evident for over-estimators. Findings suggest over- and under-estimators use differing cognitive processes when assessing their performance, such that under-estimators rely on recollection during memory and over-estimators draw upon excess familiarity when over-estimating their performance. Episodic memory thus appears to play a contributory role in metacognitive judgments of illusory superiority and inferiority.

**Graphical Abstract:** Event-related potentials (ERPS) were recorded for the Dunning-Kruger Effect as subjects made metacognitive judgments about performance on a memory task. Over- and Under-estimators exhibited a crossover interaction in response times when believing they did best and worst, respectively. A crossover pattern was also observed for ERPs: LPC signals of recollection were found for under-estimators, whereas familiarity-based FN400 effects were evident for over-estimators and correlated with estimates.

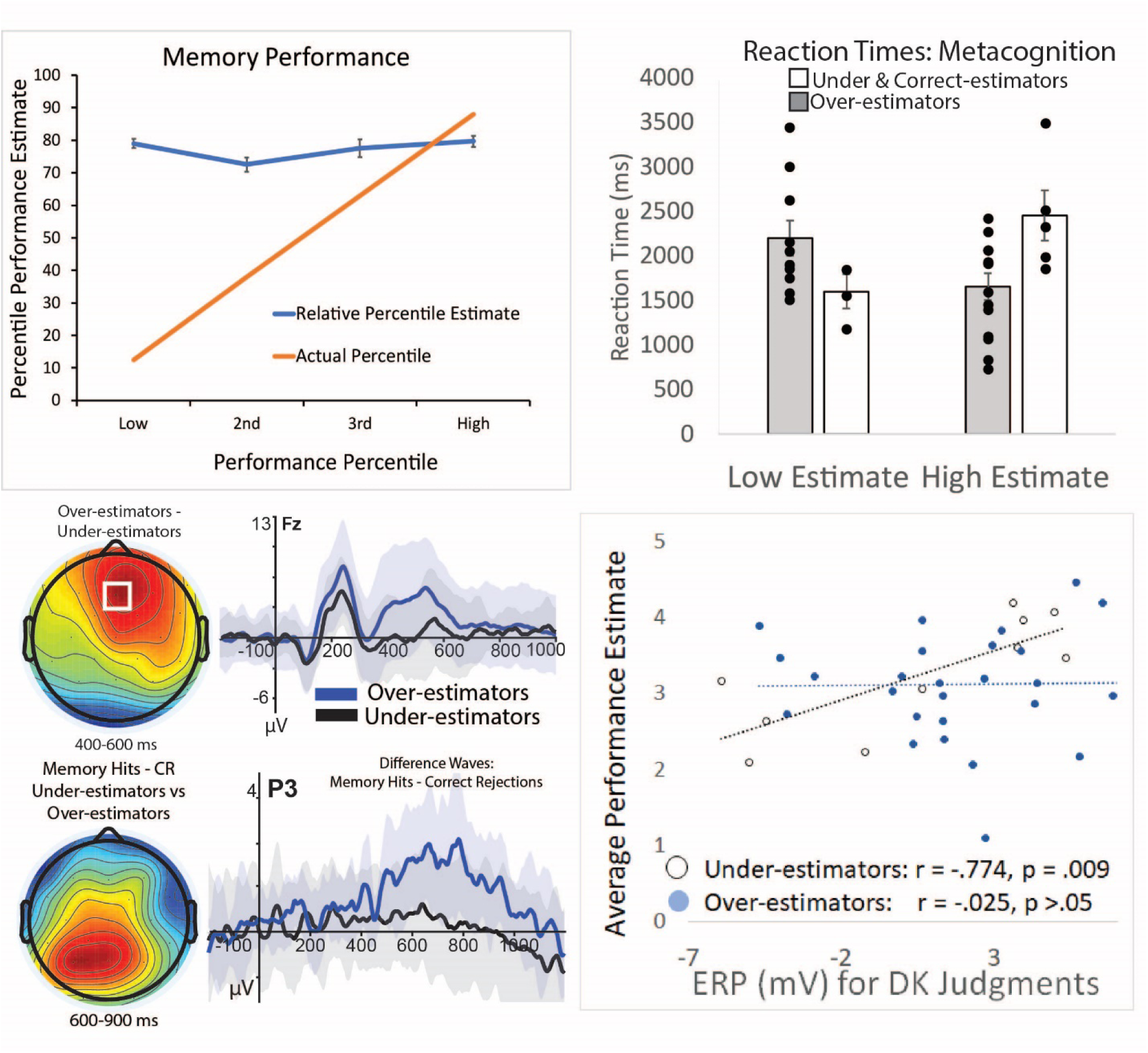

## Introduction

> “*The fool doth think he is wise, but the wise man knows himself to be a fool*” (Shakespeare, 1601)

The Dunning-Kruger Effect (DKE) describes the phenomenon in which poor performers on a task tend to overestimate their performance while high performers on a task tend to underestimate their performance. Overconfidence has been a topic of interest throughout recorded history, as early as the time of Socrates, who was noted by Plato for identifying *“…it is likely that neither of us knows anything worthwhile, but he thinks he knows something when he does not, whereas when I do not know, neither do I think I know; so I am likely to be wiser than he to this small extent, that I do not think I know what I do not know”*, and later with Charles Darwin noting more simply that “*Ignorance more frequently begets confidence than does knowledge*” (Darwin, 2009/1871). Also implied by these timeless observations is that these metacognitive illusions are bi-directional, such that the more competent individuals who perform highest tend to also under-estimate their abilities (for Review see Zell et al., 2019).

Perceptions of inaccurate confidence in one’s abilities is a common phenomenon that can happen to anyone (including the authors and readers) and can lead to serious problems that are often preventable. For example, the overconfidence that the Titanic was unsinkable led to loss of over 1500 lives (Bartlett, 2012; Lord, 1986; Lord, 1955) and in modern times, the COVID-19 pandemic of 2020 has been noted for widespread over-estimation of abilities to manage the pandemic concurrent with early under-estimation of its world-wide impact by many world health organizations, governments, and media alike (with a few exceptions). Conversely, when the most competent people hold contributions from teams or society because they think others are better suited, the effects can also lead to serious problems that can be a substantial loss on society. For example, we could lose the most competent people for leadership positions, and instead embrace those whose merely think they are the best (incorrectly). Therefore, it is important to understand how and why these inaccurate judgments of one’s abilities occur, so that they can be prevented.

In 1999, the study of over- and under-confidence was further characterized by David Dunning and Justin Kruger, social psychologists who explored combining the two effects under one term. In their landmark study, Dunning and Kruger conducted several studies showing that bottom performers on a tests of humor judgments, logical reasoning, and grammar overestimated their performance percentile and that, conversely, top performers underestimated their performance percentile. The name “The Dunning-Kruger Effect” became highly popularized throughout mainstream culture and society (Herrera, 2018; Marczyk, 2017). Generically, the DKE describes a phenomenon in which self-estimates of performance on a task and the related percentile ranking among others also participating in the task do not match actual performance. The direction of this mis-match of self-perception extends in both directions (Sieber, 1979). Empirically, this paradigm has been used successfully in many different tasks to elicit the DKE on such tasks as microeconomics college exams (Ryvkin, Krajč, & Ortmann, 2012b), knowledge about the University of Chicago (Burson et al., 2006), logical reasoning (Schlösser et al., 2013), cognitive reflection (Pennycook et al., 2017), size judgments (Sanchez, 2016), finance (Atir, Rosenzweig, & Dunning, 2015), and computer programming (Critcher & Dunning, 2009). More broadly, the effect has been referred to in popular culture contexts of aviation, (Woodbury, 2018), driving (Svenson, 1981), and professors rating their own teaching skills (Cross, 1977). However, an account of the cognitive processes leading to these illusory experiences has yet to be fully explored.

### The Dunning-Kruger Effect

The DKE is a psychological phenomenon in which a mismatch in one’s perceived ability and the reality of one’s objective performance on a given task appears to be directionally moderated by the factor of ability. Low performers (individuals who do not earn high scores on a test using an objective scale) tend to overestimate their performance percentile on a task while high performers (individuals who earn high scores measured on an objective scale) tend to underestimate their performance percentile on the same task. Most paradigms used to research the DKE follow a similar format: participants are given a task such as a series of logical reasoning problems or math problems, etc., and after they finish the task in its entirety, they are asked to estimate their overall percentile estimate and objective score on the task. That is, their metacognitive judgment is measured as a single data point assessed at the conclusion of the study and represents their aggregated assessment of performance across many trials. That approach has not, however, permitted the ability to measure multiple instances of the cognitive phenomenon per person, and has precluded being able to collect simple statistical measures of central tendency, such as reaction times. Accordingly, extant approaches have also precluded the collection of neuroscientific measures that rely upon have multiple measures per person (electroencephalography (EEG), fMRI, etc.), but which may illuminate contributory cognitive processes.

Though relatively limited in scope, researchers have investigated illusory over-confidence for decades. One of the earliest studies of overconfidence was conducted by Adams and Adams (1960) who found that participants’ confidence in their ability to recognize correctly spelled words was higher than their accuracy at the task. Five years later, Oskamp (1965) found that when clinical psychologists were asked to make a diagnosis for a case study, their confidence in their decision increased when they were given more information about the case although their accuracy did not increase. These instances showed that confidence and accuracy were not necessarily correlated, both in experimental studies and in more practical issues of clinical diagnoses. This finding has persisted in modern research on memory as well (Hirst et al., 2015; Kvavilashvili, et al., 2009) and will be discussed later below.

Memory research intersects with the DKE at the point of confidence in one’s memories and the accuracy of those memories. A large collection of research is available that supports the finding that high confidence does not beget high accuracy (Chua, et al, 2012; Roediger & DeSoto, 2014; Koriat et al., 2008; Nelson & Narens, 1990; Wells, et al., 2002). People can often exhibit illusory over-confidence in their memory judgments, and this has had wide-ranging impacts on society. For example, so-called ‘flashbulb memories’ and thought to be some of our most salient memories, yet are found to be no more accurate than other memories despite the participants’ high confidence in their accuracy (Neisser & Harsch, 1992; Brown & Kulik, 1977). Other experiments studying the ‘false fame’ phenomenon demonstrate that familiarity with names can lead to falsely recognizing them as famous later (Dywan & Jacoby, 1990; Jacoby, et al., 1989; Jacoby, Woloshyn, & Kelley, 1989, 2004): familiarity caused participants to falsely believe an ordinary name was famous because they could not recollect the context in which the name was presented. Perhaps some of the most impactful research on memory failing to correlate with accuracy pertains to the criminal justice system (Heaton-Armstrong, et al., 2006; Loftus, 1975; Loftus & Zanni, 1975; Nadel & Sinnott-Armstrong, 2012; Pena, et al., 2017; Schacter & Loftus, 2013). In a study by Pena et al. (2017) researchers asked participants to make judgments about their accuracy on a memory test for a mock crime observed earlier in a study. They found that participants who performed poorly on the memory test for details of a mock crime overestimated their memory accuracy, similar to what has been found for making prospective judgments of learning (JOL) (Irak, et al., 2019; Müller et al., 2016; Metcalfe & Finn, 2008; Koriat & Ma’ayan, 2005). These results were consistent with the results of poor performers exhibiting the DKE, demonstrating the link that may inherently exist between the two domains of memory and metacognition, yet which remains largely unexplored.

Throughout the years, overconfidence in general was studied in contexts such as social situations (Dunning, et al., 1990; Vallone, et al., 1990), tasks of differing degrees of difficulty (Bradley, 1981; Lichtenstein & Fischhoff, 1977; Sen & Boe, 1991), and ways to reduce overconfidence (Arkes, Christensen, Lai, & Blumer, 1987; Zechmeister, Rusch, & Markell, 1986). In this research, there was a common finding of people having overconfidence in wrong answers (Fischhoff, Slovic, & Lichtenstein, 1977; Harvey, 1990; Howell, 1971; Lichtenstein & Fischhoff, 1977; May, 1986) and a less common finding of under-confident correct answers for top performers (Sieber, 1979) since the focus of the research at that time was not the high performers. The term “overconfidence effect” developed to describe this pattern of higher self-estimates of confidence than ability, which stemmed into a related social psychological phenomena known as the ‘better-than-average-effect’ (BTAE), whereby people are found to rate themselves as better than average peers (for Review see Zell et al., 2019; Alicke & Govorun, 2005; Chambers & Windschitl, 2004; Moore & Healy, 2008; Sedikides, Gaertner, & Cai, 2015; Brown, 1986, 2010; Hartwig & Dunlosky, 2014). The BTAE reflects a general self-evaluation bias (Kwan et al., 2004; Moore & Healy, 2008), though as noted by Zell et al. the DKE emphasizes more of the bi-directional illusions including those believing they are less than average. Several somewhat-cognitive accounts have been put forth of these phenomena, such as ‘egocentrism’, positing that people know more about themselves than about others and hence rank themselves higher in evaluations from the richer depth of knowledge (Moore & Healy, 2008), but as noted by Zell et al., (2019) such accounts also have not been able to fully explain the underlying mechanisms of what is driving these kinds of social psychological phenomena of over-confidence in one’s abilities (for alternative views of the DKE, see: Gignac & Zajenkowski (2020), Sullivan, Ragogna, & Dithurbide (2018), Mahmood (2016), Karjc & Ortmann (2008), Kreuger & Mueller (2002)).

### Theoretical Accounts of the Dunning-Kruger Effect

Dunning and Kruger postulated that the reason for low performers’ incorrect estimation for objective performance scores is due to meta-ignorance or two-fold ignorance (Kruger & Dunning, 1999). This means that poor performers are unaware that they are ignorant of the details needed to correctly complete the task and that double ignorance bolsters feelings of false superiority (Schacter, 2012). More simply, poor performers do not have the knowledge to complete the task correctly and because they do not know their answers are incorrect, they believe they are performing well (i.e. ‘ignorance is bliss’) (Schlösser et al., 2013). While this is a very useful behavioral description, it does little to advance an understanding of the cognitive processes involved in the pervasive illusion.

Dunning and Kruger also used what they coined ‘reach-around-knowledge’ to explain low performers’ high confidence in their abilities. The term ‘reach-around knowledge’ refers to a person’s unique knowledge gained from previously participating in a similar task to the task presented and generalizing their past experiences to the current experience (Dunning, 2011). Kruger and Dunning postulated that participants use ‘reach-around-knowledge’ to help achieve their estimation, though this doesn’t necessarily require that it leads them to an accurate perception. According to this view, in order to give an overestimation, one must first have knowledge about the same or similar tasks but not have the knowledge about the details of the task to complete it correctly. Having a larger store of reach-around knowledge should therefore increase the overestimation of poor performer’s scores. On the contrary, having a smaller store of reach-around knowledge should decrease the overestimation of one’s abilities resulting in a more accurate performance estimate.

Dunning and Kruger’s ‘reach-around-knowledge’ account, however, is only a theoretical concept that has not yet been operationally defined or objectively measured, and also lacks a substantive construct grounded in cognitive psychology. Nevertheless, it provides a useful platform from which to expand upon in investigating this phenomenon within a theoretical construct. The ‘reach-around-knowledge’ account provided by Dunning and Kruger refers to changes in current behavior based upon prior experience, which is fundamentally a defining feature of memory (Rudy, 2013), and as such it recognizes a key role that memory processes may play in contributing to this metacognitive illusion. There happens to be a rich and robust empirical history of memory processes being both theoretically and operationally defined and studied. Two of these cognitive processes of episodic memory that may contribute to the DKE are familiarity and recollection (Yonelinas, 1999; 2002; Eichenbaum et al., 2008; Yonelinas et al., 2010). These processes align closely with the general concepts that Dunning and Kruger attributed to their ‘reach-around-knowledge’ account, and importantly, we can draw upon this platform of memory processes in approaching the DKE in a systematic manner, which will be discussed in depth in the sections below.

### Familiarity and Recollection

The cognitive processes of familiarity and recollection have featured prominently in theoretical models of episodic memory for several decades (Eichenbaum, Yonelinas, & Ranganath, 2007; Khoe, Kroll, Yonelinas, Dobbins, & Knight, 2000; Rugg & Curran, 2007; Squire, Wixted, & Clark, 2007; Yonelinas, 1999, 2002). Broadly, familiarity refers to having exposure to some material but not being able to recall the context in which it was presented, usually associated with mid-levels of confidence, whereas recollection refers instead to information that one can recall specific contextual details about and is typically associated only with high-confidence memories (Addante, et al., 2012a, 2012b). In explicit memory, theoretical models of recognition are largely governed by the dual processes of familiarity and recollection (Diana, Yonelinas, & Ranganath, 2008; Eichenbaum et al., 2007; Ranganath, 2010; Yonelinas, 2002; Yonelinas, Aly, Wang, & Koen, 2010) (though see Wixted, 2007 and Wixted & Mickes, 2010 for alternative views), and it is possible that understanding of familiarity and recollection processes in memory may help explain a proportion of variance in the DKE.

Recollection is typically operationalized as the declarative retrieval of episodic information of both the item and context bound together into a cohesive retrieval of the episodic event (for review see Diana et al., 2008), and in empirical studies is usually associated with the retrieval of contextual information about the item of the event, such as source memory (Addante et al. 2012a; for reviews see Eichenbaum et al., 2007; Yonelinas et al. 2010; Ranganath 2010). The item in the event, however, may be retrieved without recollection, via reliance upon familiarity, which is typically conceptualized as retrieval of an item from a prior episode but without the associated contextual information in which it occurred. Familiarity occurs, for instance, when a person can remember that someone seems recognizable from the past but cannot retrieve specific information of who the person is or from where they know them.

Recollection, on the other hand, would be remembering precisely who someone is and how you know them from a prior episode of one’s past experience. Each of these two memory phenomena have been found to be dissociable cognitive processes (Yonelinas, 2002), with dissociable neural substrates in the medial temporal lobes (Ranganath et al., 2004), neuropsychologically dissociable among patient impairments (Addante, Ranganath, Olichney, & Yonelinas, 2012; Düzel et al., 1999; Mecklinger, von Cramon, & Matthes-von Cramon, 1998; Bowles et al., 2008), and with distinct patterns of electrophysiology at the scalp that is both spatially and temporally dissociable in event-related potentials (ERPs) (Addante et al., 2012; Curran, 2000; Friedman, 2013; Gherman & Philiastides, 2015; Rugg et al., 1998; Rugg & Curran, 2007). Physiologically, familiarity has been associated with ERP differences in old and new memory trials during a negative-going peak at the mid-frontal scalp sites at approximately 400 milliseconds to 600 milliseconds post stimulus, called the mid-frontal old-new effect, or FN400 (for frontal-N400 effect). On the other hand, recollection has been associated with differences between memory conditions occurring at a peak in the ERP at the parietal region of the scalp from approximately 600 milliseconds to 900 milliseconds called the late parietal component, or LPC (Addante et al., 2012; Leynes et al., 2005; for reviews see Rugg & Curran, 2007; Friedman, 2013).

### A Memory-Based Framework of the Dunning-Kruger Effect

Many of the accounts of the DKE have focused primarily upon interpretations based upon metacognition and competency (Adams & Adams, 1960; Ehrlinger & Dunning, 2003; Kruger & Dunning, 1999; Oskamp, 1965; Pennycook et al., 2017; Ryvkin et al., 2012a; C. Sanchez & Dunning, 2017). However, it is very likely that DKEs could also be contributed to by memory experiences in one’s past influencing the real-time processing of the current information-either via explicit or implicit means. Based upon the converging literatures from memory and metacognition, a viable alternative theory we postulate to explain the DKE is that illusory superiority may also be driven, at least in part, by increased familiarity from prior experience of one’s past with the tested materials (e.g.: Chua et al., 2012). People may use a decision heuristic inducing a sense of performing well despite a lack of specific retrieval of the relevant details that would be involved in marking competency with the material. In this view, the experience of lacking a distinct recollection for but being generally familiar with material will lead people to assume that they are competent and successful at the task. This scenario would be associated with increased FN400 amplitudes in ERPs for inaccurate over-estimators.

In that case, it would be a relatively ‘dangerous’ combination to have insufficient recollection but excessive familiarity with a given topic, stimuli, or information (Chua et al., 2012) because it could lead to inaccurate over-estimates of one’s abilities and competencies. Accordingly, under-estimators may be marked by having had sufficient recollection of the studied material (e.g. competency), such that these instances are associated with an LPC, while also leading people to recollect the extent of non-criterial information that their cognizance acknowledges could still be relatively wrong (Parks & Yonelinas, 2007), hence lowering their estimated scores relative to other people. In this case, the excess of recollection signal would outweigh the noise of the familiarity signal.

### Current Study

The current paradigm has been designed to study the metacognitive decision-making process as it occurs in real-time during DKE percentile estimates that are provided by participants throughout an item recognition memory test. Based upon the memory framework for making metacognitive decisions noted above, broadly, we hypothesized that low performers who tend to overestimate their percentile ranking may do so because of familiarity for previous experiences in similar situations with markedly less reliance on recollection; and that high performers who tend to underestimate their percentile ranking may use more recollection to accurately outperform their peers. Accordingly, we hypothesized that a larger FN400 will be evident in the group level ERPs for low performers compared to high performers and that there would be a larger LPC evident in the ERPs of high performers compared to low performers.

## Methods

### Participants

The experiment was conducted as approved by the California State University – San Bernardino Institutional Review Board protocol for research on human subjects. Participants were recruited through a combination of methods including advertisements placed around CSUSB or through the school-wide research pool SONA. Participants recruited through advertisements were paid $10 an hour for sessions that lasted approximately 2 hours. The study consisted of 61 right-handed participants (48 female) who were students at California State University, San Bernardino and reported being free from neurological and memory problems. Four participants’ data were not used due to noncompliance issues (pressed only 1 button throughout the task or ignored experimenter’s instructions) and one participant did not have usable data due to a experimenter error that resulted in the loss of that data. Two participants did not have usable EEG data due to excess motion artifacts/noise that resulted in a majority of trials being excluded from EEG but were included in behavioral analyses. This presented a behavioral data set of N = 56 and EEG data set of N = 54. 56.5% of the participants were self-reported to be Hispanic, 22.6% Caucasian, 11.3% Asian, and 9.7% identified as more than one or another ethnicity. The average age was 23.52 years old (*SD* = 4.82). None of the participants reported any visual, medical, or physical issues that would interfere with the experiment. Most participants spoke English as their first language (N = 47) and the 15 whom indicated speaking a different language first had been speaking English for an average of 16.73 years (*SD* = 4.74).

### Procedure

Participants arrived at the lab and completed informed consent and demographic information forms via voluntary self-report. The paradigm used was a modified item recognition confidence test, which built from similar paradigms successfully used in our lab’s prior research (Addante et al., 2012; Addante, Watrous, Yonelinas, Ekstrom, & Ranganath, 2011; Addante, 2015; Addante, Ranganath, Olichney, & Yonelinas, 2012; Roberts et al., 2018) and described in further detail below (Fig. 1). This paradigm consisted of an encoding phase containing four study sessions, in which participants studied 54 words in each session, and a retrieval phase, containing six test sessions in which the participant’s memory was tested for 54 words in each session. They viewed a total of 324 words, 216 of which were presented in the encoding phase and 116 of which were unstudied items (new items).

**Figure 1.**
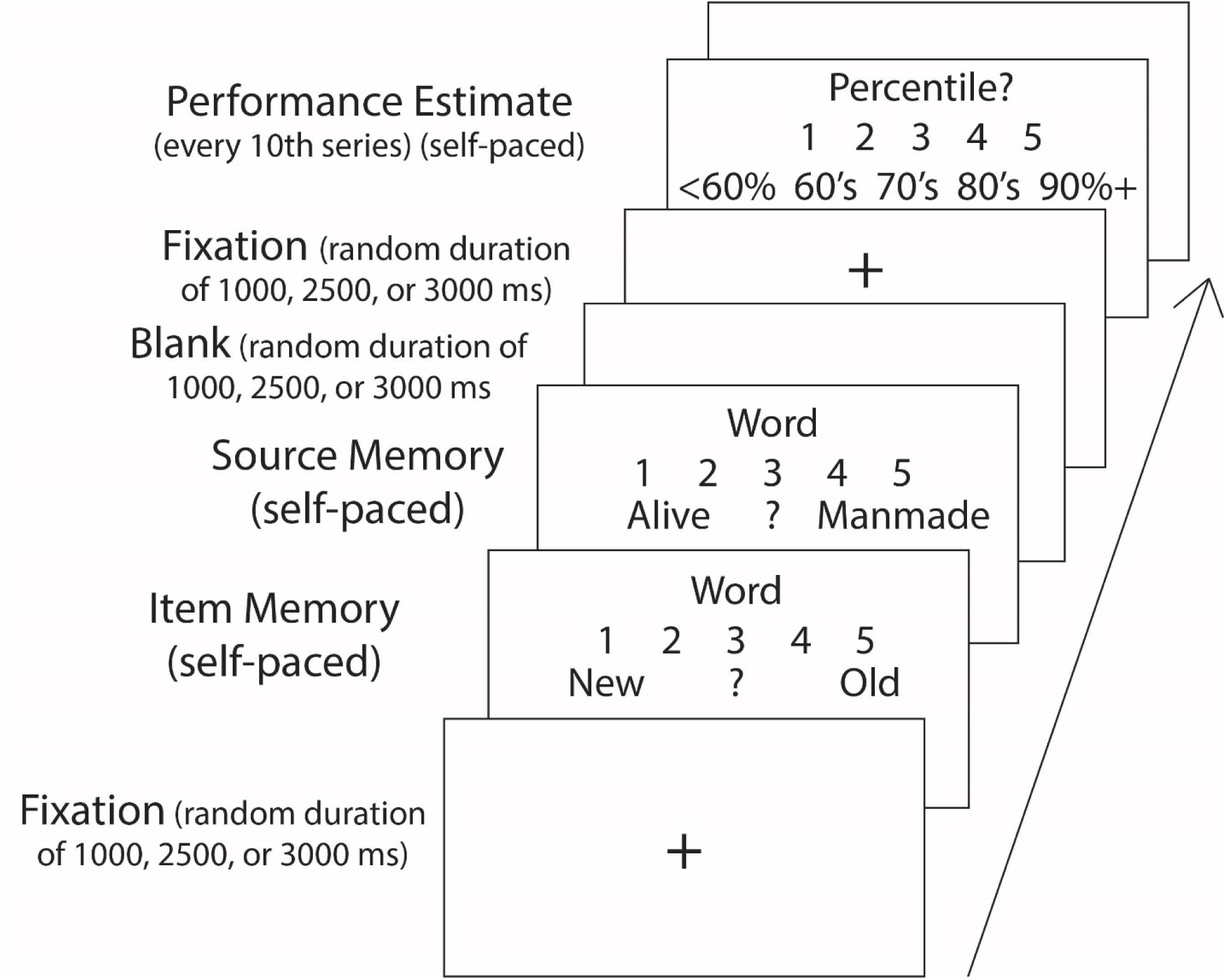
Memory Retrieval and Dunning-Kruger Testing Paradigm. Left: Participants indicated their confidence for item memory and source memory. For every 10^th^ stimuli presented, the participants viewed the Dunning-Kruger Estimate: asking participants to estimate the percentile in which they believe they are performing up to that point on the task in relation to other students.

During encoding, participants were given instructions to make a simple decision about the word presented on the screen. Subjects were asked to either judge if the item was manmade or if the item was alive and conditions were counterbalanced. The stimuli were presented on a black computer screen in white letters. To begin a trial, a screen with a small white cross at the center was presented for one of three randomly chosen inter-stimuli-interval (ISI) times: 1 second, 2.5 seconds, or 3 seconds. Then, the stimulus word appeared in the middle of the screen with ‘YES’ presented to the bottom left of the word and ‘NO’ presented to the bottom right of the word. The participants indicated their answer by pressing buttons corresponding to ‘YES’ and ‘NO’ with their index and middle fingers, respectively, and this response was self-paced by the participant. After the participants responded, they viewed a blank black screen at a random duration of 1 second, 2.5 seconds, or 3 seconds. After the blank screen, the small white cross appeared at the center of the screen to begin the next trial. This cycle continued until all 54 words in all four lists were presented. Between each list, participants were read the instructions for the next task to prevent carryover effects of the preceding encoding task, and to ensure they correctly switched between the animacy and the manmade decision task.

After the encoding phase was complete, the EEG cap was sized and ocular electrodes were attached. EEG was recorded using the actiCHamp EEG Recording System with a 32-channel electrode cap conforming to the standard International 10–20 System of electrode locations. Each subject was tested individually inside a sound-attenuating chamber. Stimulus presentation and behavioral response monitoring were controlled using Presentation software on a Windows PC. EEG was acquired at a rate of 1024 Hz. Subjects were instructed to minimize jaw and muscle tension, eye movements, and blinking. EOG was monitored in the horizontal and vertical directions, and this data was used to eliminate trials contaminated by blinks, eye-movements, or other related artifacts. Five ocular electrodes were applied to the face to record electrooculograms (EOG): two above and below the left eye in line with the pupil to record electrical activity from vertical eye movements, two on each temple to record electrical activity from horizontal eye movements, and one in the center of the forehead just above the eyebrows as a reference electrode. The EEG cap was placed on the participant’s head and prepared for electrical recording. Gel was applied to each cap site and impedances were lowered below 15 KOhms via gentle abrasion to allow the electrodes to obtain a clear electrical signal.

After the EEG cap was in place, the participant began the retrieval phase. The participants were read instructions asking them to judge if the stimulus word presented was old (studied during the encoding phase) or new (not studied in the encoding phase) (Figure 1). As in the encoding phase, all stimuli words were presented in white font on a black screen. At the beginning of each trial, participants were presented with a word in the middle of the screen, the numbers “1”, “2”, “3”, “4”, and “5” evenly spaced beneath the word, the word “New” on the left by the number “1”, and the word “Old” on the right under the number “5”. Participants pressed any number between “1” and “5” to indicate if they confidently believed the word was old (“5”), believe the word was old but was not confident (“4”), did not know if the word was old or new (“3”), believe the word was new but was not confident (“2”), or confidently believed the word was new (“1”) (Addante et al., 2011; 2012a; 2012b). This prompt was subject-paced. Participants were told to choose the response that gave us the most accurate reflection of their memory, and to respond as quickly and accurately as possible. Immediately after the item recognition judgment, participants were asked to answer a source memory confidence test indicating if the word came from the animacy decision task or the manmade decision task during encoding on a scale of 1 to 5, which was also subject-paced (Addante et al., 2011; 2012a; 2012b; 2015; Roberts et al., 2018). After responding, participants viewed a blank black screen at a random duration of 1 second, 2.5 seconds, or 3 seconds. Participants were instructed to blink only during this blank screen and avoid blinking during the screens with a small cross or stimuli.

### Dunning-Kruger In-test Questions

After the source memory test for each 10^th^ word presented during the memory test, the Dunning-Kruger estimate was presented. Participants received instructions asking them to estimate the percentile in which they believed they were performing up to that point in the test compared to other students who would participate in the study (subjects were instructed to focus on generic memory performance and to use their item memory as the primary context). During the test phase, the word “Percentile?” was presented as a prompt for their estimate with the numbers “<60%,”, “60’s”, “70’s”, “80’s”, and “90%+” evenly spaced beneath it. The Dunning-Kruger estimate was subject-paced.

### Dunning-Kruger Post-Test Questions

At the conclusion of the memory retrieval test, participants answered four post-test questions First, they were asked to “Estimate your score on the whole test”. Participants were prompted to respond on a 5-point scale with “1” meaning below 60%, “2” meaning between 60 and 69%, “3” meaning between 70 and 79%, “4” meaning between 80 and 89 percent, and “5” meaning above 90%. The second question they were asked was the following: “In what percentile did you perform on the whole test?”. The participants were prompted to respond on a 5-point scale with “1” meaning below the 60^th^ percentile, “2” meaning between the 60^th^ and 69^th^ percentile, “3” meaning between the 70^th^ and 79^th^ percentile, “4” meaning between 80^th^ and 89^th^ percentile, and “5” meaning in the 90^th^ percentile or above. The first questions measured perceived objective score on the entire memory test while the second question measured perceived relative score in relation to other students taking the memory test. These post-test prompts allowed us to test for the DKE at a between-subjects level to be sure the effect can be elicited using an episodic memory task. During analyses, participants were grouped into quartiles based on their percentile score on the test, allowing us to average each group’s responses and test them against the other group’s average responses to determine significant differences. They were also grouped by errors in percentile estimates; groups of over-estimators, correct-estimators, and under-estimators (also referred to as estimator groups later) were also made to investigate potential differences in cognitive strategies (see below).

The two additional post-test questions were: 1) “Rate your memory in everyday life” and 2) “How difficult was this entire test?”. For the first question, participants responded on a 5-point scale with “1” meaning very poor, “2” meaning poor, “3” meaning moderate, “4” meaning good, and “5” meaning very good. For the second prompt, participants responded on a 5-point scale with “1” meaning very hard, “2” meaning hard, “3” meaning moderate, “4” meaning easy, and “5” meaning very easy.

### Dunning-Kruger Groupings

To maintain consistency towards replicating the original report by Kruger and Dunning (1999), subjects were grouped for analyses in the same fashion as the original paper by separating participants into four quartiles depending on their test accuracy and investigating group differences among those quartiles. The way that subjects were selected for their respective group membership was based upon their performance on the item recognition test, divided into four quartiles. Subjects’ accuracy scores on the memory task (measured as the probability of a hit minus the probability of a false alarm, pHit-pFA) were ranked from smallest to largest and split into quartiles of performance (<=25%, >25% to 50%, >50% to 75%, and >75%) and participants who fell into these quartiles comprised the low, 2^nd^, 3^rd^, and highest quartile.

Participants were then regrouped into what we call ‘estimator groups’ or participants that over-estimated, correctly estimated, and under-estimated their percentile ranking. To make these estimator groups, first, the percentile rankings described above were given scores of 1 through 5 that directly corresponded to the scale used by subjects to estimate their performance percentile group both in- and post-test. For example, a participant who scored in the 21^st^ percentile was assigned a value of 1 while a participant who scored in the 82^nd^ percentile was assigned a value of 4.

This allowed the subtracting of each actual percentile score from the participant’s estimated percentile score (estimated percentile – actual percentile) on the post-test measure^1^, thus obtaining a value of how accurately participants estimated their percentile ranking. Positive values indicated over-estimations, values of 0 indicated correct estimations, and negative values indicated under-estimations. As an example, a participant who estimated their score to be in the range of the 80^th^-89^th^ percentile (which would correspond to a response of 4 on the response scale) and yet actually performed in the 74^th^ percentile (a corresponding response of 3) would be categorized as an over-estimator. Thus, these new groups became our over-estimators (N = 38), correct estimators (N = 8), and under-estimators (N = 10). From these group memberships, the ERP analyses were conducted in accordance with standard practices in the field of including only subjects who met a sufficient number of trials in each condition of the ERP comparison for signal-to-noise ratio (see Methods section below for full details) (over-estimators (N = 36), correct estimators (N = 8), and under-estimators (N = 10).

### Electrophysiological Analyses

Physiological measurements of brain activity were recorded using EEG equipment from Brain Vision, LLC. All EEG data was processed using the ERPLAB toolbox using Matlab (Delorme & Makeig, 2004; Lopez-Calderon & Luck, 2014). The EEG data was first re-referenced to the average of the mastoid electrodes, passed through a high-pass filter at 0.1 hertz (Hz), and then down-sampled to 256 Hz. The EEG data was epoched from 200 milliseconds prior to the onset of the stimulus to 1200 milliseconds after the stimulus was presented, and then categorized based on performance and response accuracy.

Independent components analysis (ICA) was performed using InfoMax techniques in EEGLab (Bell & Sejnowski, 1995) to accomplish artifact correction and then resulting data was individually inspected for artifacts, rejecting trials for eye blinks and other aberrant electrode activity. During ERP averaging, trials exceeding ERP amplitudes of +/-250 mV were excluded. Using the ERPLAB toolbox (Lopez-Calderon & Luck, 2014), automatic artifact detection for epoched data was also used to identify trials exceeding specified voltages, in a series of sequential steps as noted below.

Simple Voltage Threshold identified and removed any voltage below −100 ms. The Step-Like Artifact function identified and removed changes of voltage exceeding a specified voltage (100 uV in this case) within a specified window (200 ms), which are characteristic of blinks and saccades. The Moving Window Peak-to-Peak function is commonly used to identify blinks by finding the difference in amplitude between the most negative and most positive points in the defined window (200 ms) and compared the difference to a specified criterion (100 uV). The Blocking and Flatline function identified periods in which the voltage does not change amplitude within the time window. An automatic blink analysis, Blink Rejection (alpha version), used a normalized cross-covariance threshold of 0.7 and a blink width of 400 ms to identify and remove blinks (Luck, 2014).

In order to maintain sufficient signal-to-noise ratio (SNR), all comparisons relied upon including only those subjects whom met a criterion of having a minimum number of 12 artifact-free ERP trials per condition being contrasted (Addante, Ranganath, & Yonelinas, 2012; Gruber & Otten, 2010; Kim et al., 2009; Otten et al., 2006; c.f. Luck, 2016). ERPs of individual subjects were combined to create a grand average, and mean amplitudes were extracted for statistical analyses. Topographic maps of scalp activity were created to assess the spatial distribution of effects. For ERP figures, a 30 Hz low pass filter was applied to ERPs so as to parallel the similar ‘smoothing’ function that ensues from taking the mean voltage between two latencies during standard statistical analyses (i.e. Addante, 2015). ERP results are reported for representative electrode sites but were also found to be reliable at surrounding 3-site clusters of electrodes unless otherwise noted.

### Behavioral Results

Recognition memory response distributions for recognition of old and new items are displayed in Table 1. Item recognition accuracy was calculated as the proportion of hits (*M* = .81, *SD* = .11) minus the proportion of false alarms (*M* = .24, *SD* = .14) (i.e. pHit-pFA) (Addante et al., 2011; 2012a, b). Participants performed item recognition at relatively high levels (Max = .87, Min = .18, *M* = .57, *SD* = .15) which was greater than chance probability, *t*(55) = 3.59, *p* < .001. In addition, participants’ accuracy for high confidence item recognition trials (‘5’s’) was significantly greater than low confidence item recognition trials (‘4’s’), *t*(55) = 9.04, *p* < .001. Source memory response distributions for recognition of old and new items are displayed in Table 2. Source memory accuracy values were collapsed to include high and low source confidence responses which were then divided by the sum of items receiving a correct and incorrect source response to calculate the proportion (Addante et al., 2012a, 2012b; Roberts et al., 2018). Mean accuracy for source memory was .30 (*SD* = .19) and was reliably greater than chance, *t*(55) = 11.78, *p* < .001. The results of item memory confidence and source memory confidence scores and ERPs replicated the previous findings of Addante, Ranganath, and Yonelinas (2012), as reported in further detail by Muller (2019).

**Table 1.** Distribution of Responses for Each Item Response as a Proportion of All <u>Memory Responses</u>

| Item Recognition Confidence | 1 | 2 | 3 | 4 | 5 |
| --- | --- | --- | --- | --- | --- |
| All Old Items | .09 | .07 | .04 | .21 | .60 |
| All New Items | .43 | .23 | .10 | .15 | .08 |
| Animacy Task | .13 | .06 | .04 | .16 | .60 |
| Manmade Task | .08 | .04 | .03 | .14 | .71 |

**Table 2.**
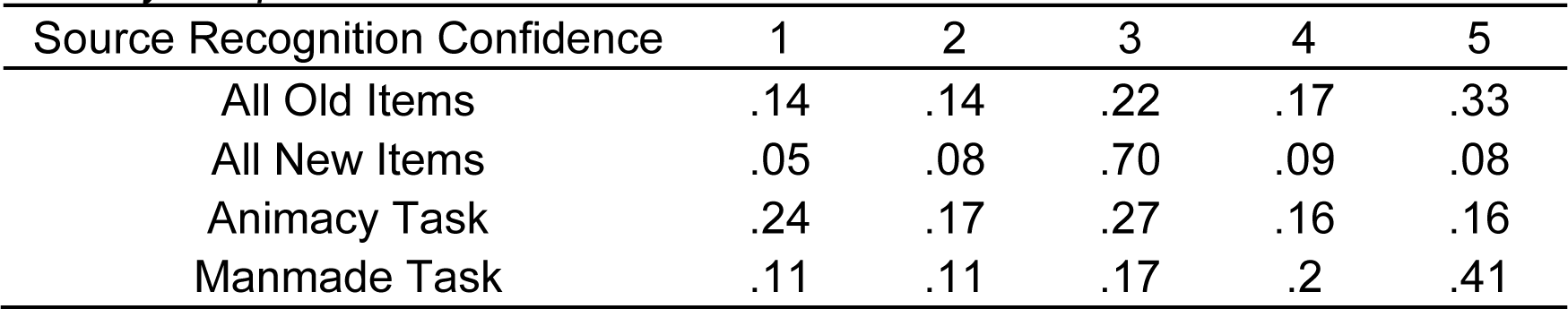
Distribution of Responses for Each Source Response as a Proportion of All Memory Responses

| Source Recognition Confidence | 1 | 2 | 3 | 4 | 5 |
| --- | --- | --- | --- | --- | --- |
| All Old Items | .14 | .14 | .22 | .17 | .33 |
| All New Items | .05 | .08 | .70 | .09 | .08 |
| Animacy Task | .24 | .17 | .27 | .16 | .16 |
| Manmade Task | .11 | .11 | .17 | .2 | .41 |

### Dunning-Kruger Performance Judgments

The distribution of responses for each Dunning-Kruger response category for the post-test and in-test Dunning-Kruger responses are shown in Table 3. When plotted against actual performance, results from subjects’ reported performance estimates revealed that the canonical DKE was evident, thereby replicating the DKE and extending it to our novel episodic memory paradigm (Figure 2). To quantify and analyze this effect, the participants were first split into quartiles based on memory accuracy (the procedure for grouping of subjects into groups based upon estimated performance vs actual performance was described in detail earlier in the Methods). The average memory test accuracy, organized by quartile and each quartile’s respective average post-test Dunning-Kruger response, is listed in Table 3.

**Table 3.**
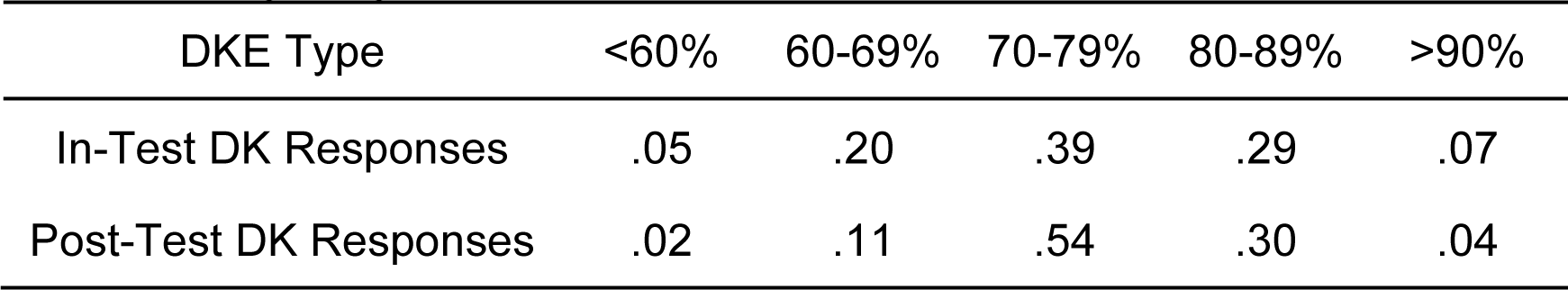
Distribution of Responses for Each Dunning-Kruger Response, as a Proportion of All Dunning-Kruger Responses

| DKE Type | <60% | 60-69% | 70-79% | 80-89% | >90% |
| --- | --- | --- | --- | --- | --- |
| In-Test DK Responses | .05 | .20 | .39 | .29 | .07 |
| Post-Test DK Responses | .02 | .11 | .54 | .30 | .04 |

**Figure 2.**
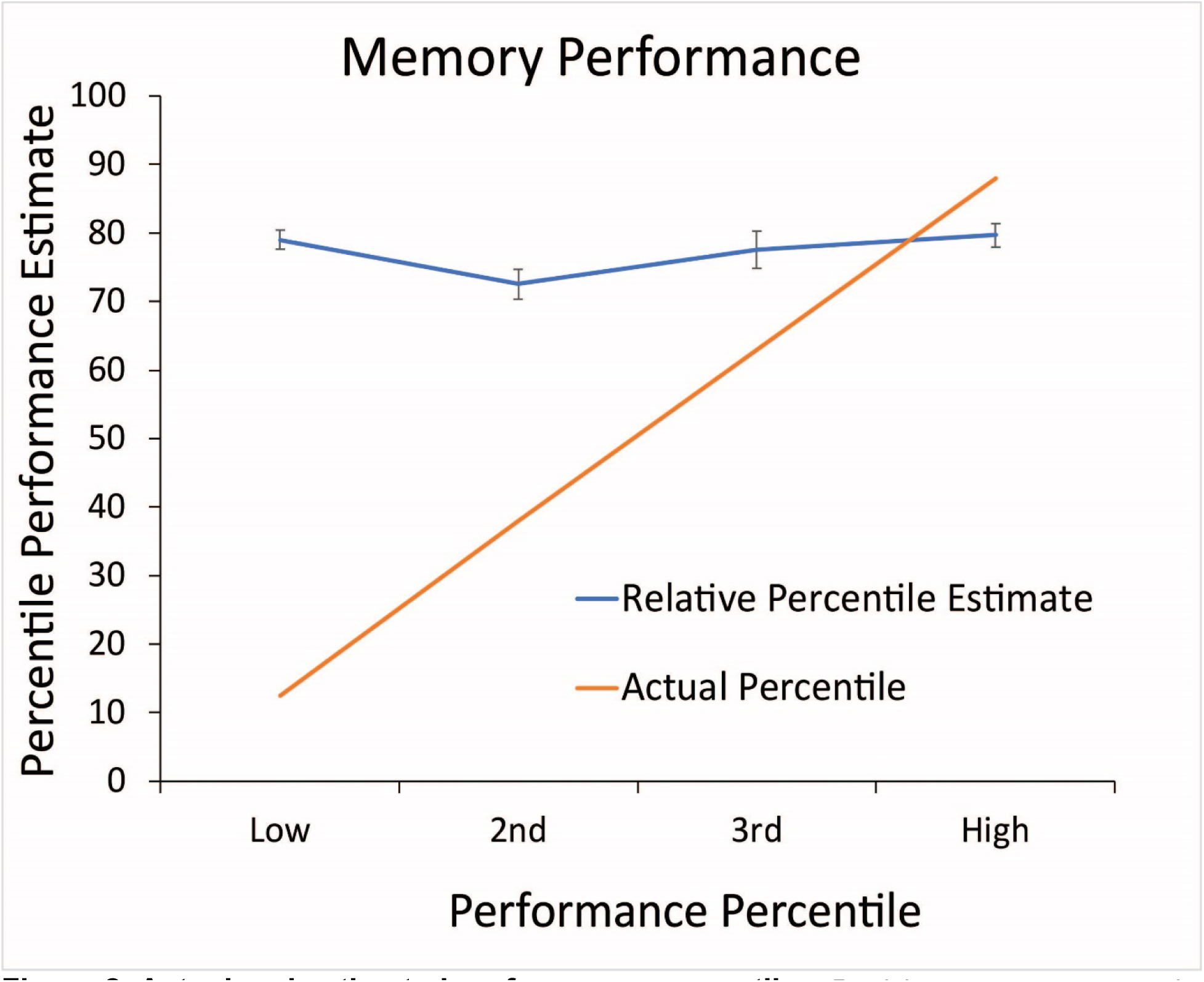
Actual and estimated performance percentiles. Participants were separated by their actual percentile ranking. The low group consists of those in the first quartile (less than or equal to 25%), the second group consists of those in the second quartile (>25% and <=50%), the third group consists of those in the third quartile (>50% and <=75%), and the high group consists of those in fourth quartile (>75%). Participants who performed in the first quartile showed the most overestimation while participants who performed in the fourth quartile showed underestimation of their actual percentile.

The bottom quartile (N = 14, *M* = 2.43, *SD* = 0.51, *t*(26) = 17.69, *p* < .001), 2nd quartile (N = 14, *M* = 1.79, *SD* = 0.80, *t*(26) = 8.33, *p* < .001), and 3rd quartile (N = 14, *M* = 1.43, *SD* = 1.28, *t*(26) = 4.16, *p* < .001) significantly overestimated their percentile ranking while the top quartile significantly underestimated their percentile ranking (N = 14, *M* = −0.79, *SD* = 0.89, *t*(26) = −3.29, *p* = .003) (Table 4).. Furthermore, the magnitude of the errors made by each group decreased systematically as their percentile group increased: the bottom quartile overestimated their percentile by 62.56%, the 2nd quartile overestimated by 37.95%, the 3rd quartile overestimated by 14.56%, and the top quartile underestimated by 8.30%. Together, these basic findings provide evidence that the DKE was successfully elicited by our memory paradigm, one that to our knowledge has not been shown before, thereby extending the DKE phenomenon directly to episodic memory.

**Table 4.** Average Recognition Memory Test Accuracy and Average Post-Test and In-Test Dunning-Kruger Relative Response by Quartile

| Quartile | Accuracy | Average Post-Test<br>DK Relative<br>Response | Average In-Test<br>DK Relative<br>Response |
| --- | --- | --- | --- |
| Top (N = 14) | .74 (.06) | 3.50 (0.65) | 3.26 (0.73) |
| 3rd (N = 14) | .62 (.02) | 3.29 (0.99) | 3.33 (1.01) |
| 2nd (N = 14) | .55 (.04) | 2.79 (0.80) | 2.79 (0.81) |
| Bottom (N = 14) | .38 (.08) | 3.43 (0.51) | 3.17 (0.62) |

*Note.* Standard deviations are in parentheses.

Figure 3 displays the raw performance of each subject in item recognition as a function of Dunning-Kruger groupings for their estimates of the percentile group in which they thought they were performing (over-estimator, under-estimator, or correct-estimator). Performance of the over-estimators on the item recognition memory task (N = 38, *M* = .501 *SD* = .11, *SE* = .02) was significantly less than the correct-estimators (N = 8, *M* = .65, *SE* = .02) (*t*(44) = 3.71, *p* <.001) and under-estimators (N = 10, M =.75, SE = .02) (*t*(46) = 6.66, *p* <.001), though was reliably greater than chance-level performance (*t*(37) = 27.341, *p* < .001). The worst-performing subjects (over-estimators) performed otherwise normatively, above the impaired levels of hippocampal amnesia patients (M = .30) that we have reported in previous work using the same paradigm (Addante et al., 2012b) and were otherwise consistent with normative performance of healthy adults in published prior studies using the same paradigm (Addante et al, 2012a; Addante et al., 2011; Roberts et al., 2018).

### Parameter estimates of decision processes during memory retrieval

Another possible account of the DKE results is that subjects may have differentially engaged with the task, or that results may reflect different decision-making strategies (we thank two anonymous Reviewers for raising these possibilities). To assess this possibility, we conducted analyses to quantify if subjects were using any discernably different decision processes reflecting differential engagement in the memory task, using the ROC Toolbox (Koen et al., 2017) to calculate parameter estimates of their Decision Criterion (C), Response Bias (B), Recollection (R_o_), and Familiarity (F) process contributions to performing the memory task (Yonelinas, 2002; 2004; Park & Yonelinas, 2011; Yonelinas et al., 2010; Koen et al., 2017). A one-way ANOVA was performed to identify potential differences among groups (under-estimators, correct-estimators, and over-estimators) on each of the parameters. There were no reliable differences observed among groups for Recollection estimates (*F*(2,52) = .75, *p* = .48), Decision Criterion (*F*(2,52) = 1, *p* = .38), or Response Bias (*F*(2,52) = .32, *p* = .73) (Table 6). Furthermore, a 3 x 5 ANOVA using factors of Group (over-, under- and correct-estimators) and Response (DK judgments of 1, 2, 3, 4, & 5) revealed no differences in how subjects distributed their metacognitive judgments across the scale (F(8,265) = 0.58, p = .79), indicating no evident differences in how groups of subjects were responding to the task.

Analyses of groups did reveal a significant effect for the parameter estimates of familiarity (*F*(50,2) = 14.35, *p* < .001) (one subject of the over-estimating group was removed as an outlier for exceeding three standard deviations from the mean) consistent with the groups’ defined memory differences noted earlier (Figure 3). This effect was further explored with follow-up between-group t-tests, which revealed that each group was reliably different from each other in their estimates of familiarity used during item recognition: Under-estimators (N = 8, *M* = 1.70, *SD* = .45, *SE* = .14) were greater than Correct-estimators (N = 10, *M* = 1.30, *SD* = .28, *SE* = .10) (*t*(16) = 2.22, *p* = .041); Correct-estimators were greater than Over-estimators (N = 35, *M* = .89, *SD* = .46, *SE* = .08) (*t*(41) = 2.389, *p* = .022), and Under-estimators were greater than Over-estimators (*t*(43) = 4.96, *p* = <.001) (Figure 3), corresponding to the underlying performance differences on the memory test between groups. Thus, outside of their core performance on the task, it appears that subjects were meaningfully engaged in the task in similar ways unattributable to factors of strategies, task-engagement, or decision-making differences of the groups.

**Figure 3.**
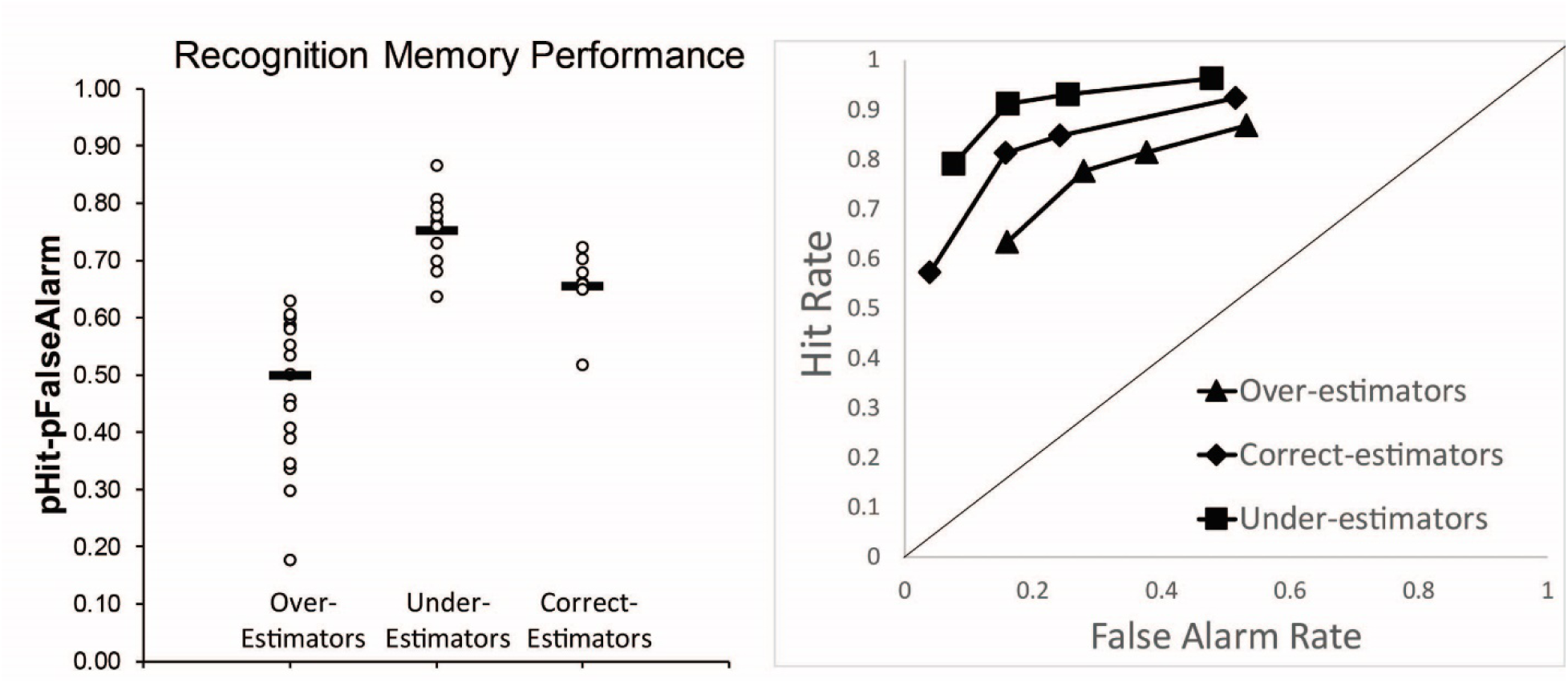
Recognition Memory Performance per Estimation Group. Left: Accuracy on Item Recognition Memory Test for Over-estimators, Under-estimators, and Correct-Estimators. Raw scores of each subject are shown plotted as grouped by estimates of performance percentile relative to the group. Memory accuracy was measured as probability of a hit minus the probability of a false alarm, plotted on the y-axis; black bars represent mean values per group. Right: Receiver operating characteristic curve of item recognition memory performance for each of the three groups. Y-axis plots the proportion of hits, x-axis plots the proportion of false-alarms for each level of confidence.

### Response Speed for Dunning-Kruger Judgments

Reaction times (RTs) were unable to be measured in previous studies of the DKE because extant studies were limited by including only a single measure of self-estimate of performance at the end of a task, which is not adequate for RT analyses. The current study, however, collected thirty DKE judgements per subject (n = 30), which permitted analysis of response times during these phenomena, in order to gain insight into how the different groups might have performed the task differently. (Table 5). A one-way ANOVA first compared general RTs collapsed across all DKE metacognitive responses (‘1’ through ‘5’) between the three groups of over-estimators (N = 38), under-estimators (N = 10), and correct-estimators (N = 8), revealing no significant differences in overall response times among groups (*F*(2, 52) = .41, p = 0.666).

However, because our hypotheses were specifically interested in how people made illusory metacognitive judgments of being either among the best or the worst performers, we also specifically analyzed of the reaction times for when subjects reported performing either the best (‘5’) or the worst (‘1’), as a function of estimator-group to explore potential differences in cognitive strategies used to make these illusory self-estimates. For this analysis, the under-estimator group (N = 10) is inherently defined as having a limited number of trials of responding that they believed they were the best percentile, and so this naturally reduced the sample of available subjects with sufficient trials for these sensitive behavioral analyses (N = 3). Although the current paradigm has been previously established as being sensitive to small samples of the same sizes (Addante et al, 2012; Addante, 2015), we nevertheless sought to increase the sample size of those who did not exhibit the errors of illusory superiority. Therefore, we collapsed the under-estimator group (N = 3) together with the available subjects of correct-estimators whom had responses in these otherwise-rare categories (N = 5) to create a larger group for analyses (N = 8); the over-estimators with available trials in these conditions were N = 23. We used a two-factor between-subjects ANOVA to determine if any mean differences existed among the reaction times for Dunning-Kruger groups (over-estimators vs. correct and under-estimators) while they judged themselves to be in the highest (response of ‘5’) and lowest (response of ‘1’) performance groups, respectively. The ANOVA revealed a significant condition by group interaction between Dunning-Kruger groups and responses, *F*(1,27) = 8.35, *p* = .008, which we explored with planned t-tests.

Within-group, the reaction times for over-estimators when rating themselves as performing in the worst (< 60^th^) percentile (DK response of ‘1’; *M* = 2204 ms, *SD* = 628 ms, N = 10) were significantly slower than when over-estimators rated themselves in as being in the best 90th percentile (DK response of ‘5’; *M* = 1656 ms, *SD* = 544 ms, N = 13), *t*(21) = 2.24, *p* = .04. Alternatively, the combined group of correct + under-estimators exhibited RTs with the opposite pattern: showing a slower response time when rating themselves in the 90th percentile or above (*M* = 2457 ms, *SD* = 634 ms, N = 5) that was marginally faster when rating themselves as performing less than the 60th percentile (*M* = 1604 ms, *SD* = 329 ms, N = 3; *t*(6) = −2.12, *p* = .08) (Figure 4).

**Figure 4.**
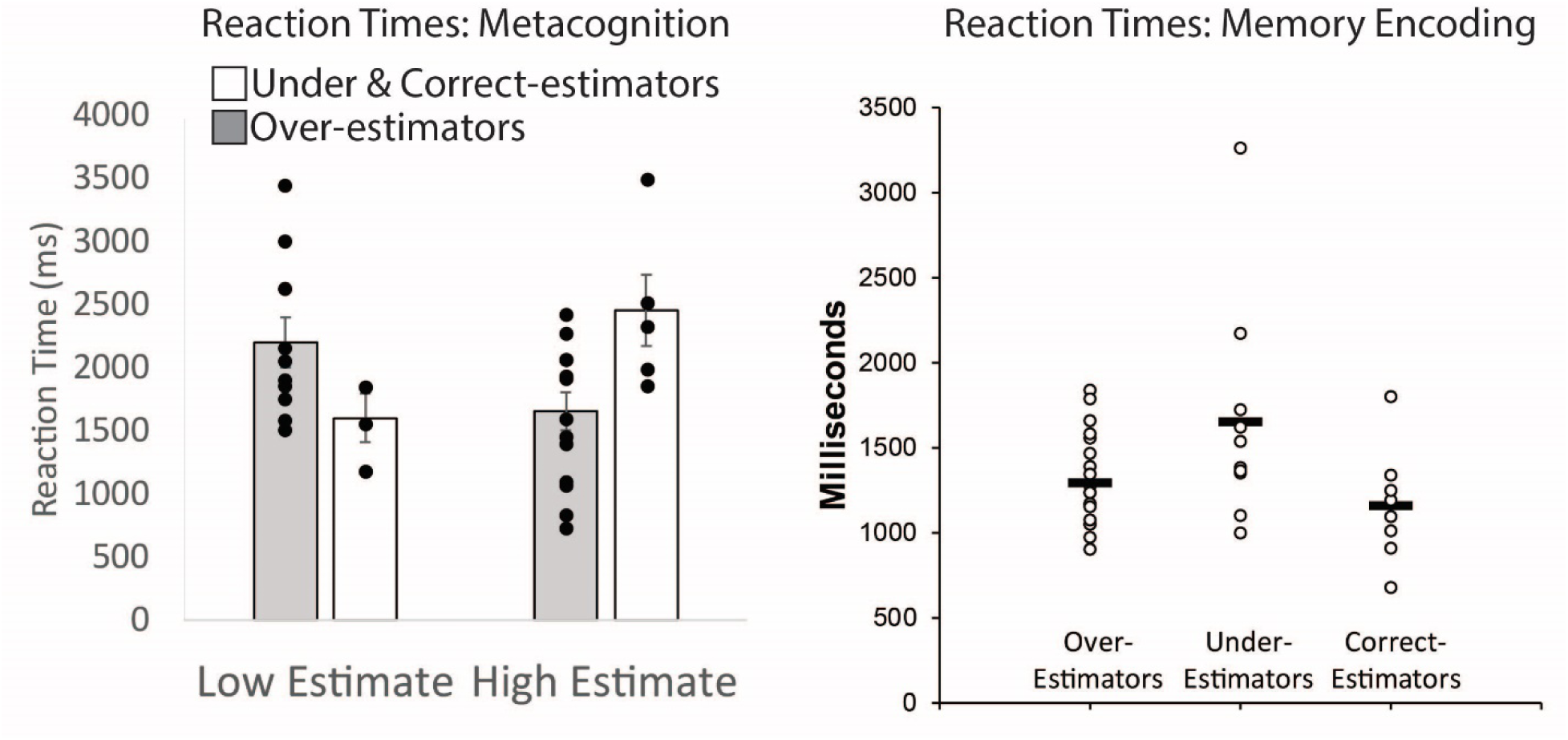
Reaction Times. Left: Mean reaction times of high and low percentile estimation by Dunning-Kruger groups. Participants’ belief of performing in the 59^th^ percentile or below corresponds to response of ‘1’ on the task and performing in the 90^th^ percentile or above corresponds to response of ‘5’. The reaction times are separated by over-estimators and the combined group of correct-& under-estimators collapsed due to relatively small sample sizes individually for these response bins. Mean reaction times are reported in milliseconds. Each black dot represents the raw score of an individual subject for each respective condition. Error bars represent standard error of the mean. Right: reaction times for the memory encoding task per each group. Black bar represents the mean for each group.

Between-groups, over-estimators were significantly faster (*M* = 1656 ms, *SD* = 544 ms, N = 13) than the collapsed group of accurate- and under-estimators (*M* = 2457 ms, *SD* = 635 ms, N = 5); when each were responding that they thought they were doing the best (i.e. in the 90^th^ percentile or above), *t*(16) = −2.68, *p* = .02). The converse pattern was evident when rating themselves in the 59^th^ percentile and lower (Table 5), as the over-estimators responded relatively more slowly (*M* = 2204 ms, *SD* = 628 ms, N = 10) than the combined group of correct- and under-estimators (*M* = 1604 ms, *SD* = 330 ms, N = 3), though this general pattern was found not to be significant for the current sample sizes, *t*(11) = 1.56, *p* = .15. This was an exploratory analysis using small samples that, like all scientific findings, will benefit from corroboration by independent laboratories. However, these results persisted despite the small sample sizes of the groups, and the patterns suggest that future studies using larger groups may find similar patterns. The pattern of responding revealed evidence that people who erred to over-estimate their abilities were also responding faster when they believed they were doing the best and slower to say they were the worst, whereas more-accurate under-estimators were slower to say they were the best and relatively quicker to say they were the worst.

**Table 5.** Response Distribution Proportions of Dunning-Kruger Responses and Mean Reaction Times, Standard Deviations, and Sample Size for In-Test Dunning-Kruger Judgments Organized by Estimator Group.

| Group | Dunning-Kruger Judgments | 1 | 2 | 3 | 4 | 5 |
| --- | --- | --- | --- | --- | --- | --- |
| Over-Estimators<br>(n = 36) | Response Distribution | .05 | .19 | .39 | .28 | .09 |
|  | Reaction Time | 2204 | 2064 | 1948 | 2044 | 1656 |
|  | SD | 628 | 641 | 644 | 860 | 544 |
|  | N per Response | 10 | 23 | 33 | 27 | 13 |
| Correct-Estimators<br>(n = 8) | Response Distribution | .09 | .28 | .33 | .25 | .05 |
|  | Reaction Time | 1447 | 2323 | 2018 | 1920 | 2275 |
|  | SD | 263 | 987 | 890 | 733 | 360 |
|  | N per Response | 2 | 6 | 7 | 5 | 2 |
| Under-Estimators<br>(n = 10) | Response Distribution | .01 | .21 | .35 | .38 | .05 |
|  | Reaction Time | 1918 | 2074 | 2166 | 1996 | 2579 |
|  | SD | -- | 1249 | 543 | 770 | 478 |
|  | N per Response | 1 | 5 | 9 | 9 | 3 |
| Combined<br>Correct- and<br>Under-Estimators<br>(n = 18) | Response Distribution | 0.04 | 0.24 | 0.34 | 0.32 | 0.05 |
|  | Reaction Time | 1604 | 2209 | 2101 | 1969 | 2457 |
|  | SD | 330 | 1062 | 693 | 729 | 635 |
|  | N per Response | 3 | 11 | 16 | 14 | 5 |

*Note.* Means and SD are in milliseconds.

**Table 6.** Memory Performance Parameter Estimates.

| DK Group | Recollection Estimate (Ro) | Familiarity Estimate (F) | Criterion (C) | Bias (B) |
| --- | --- | --- | --- | --- |
| Over-estimators (N=36) | .37 (.23) | .73 (1.08) | -.08 (.42) | 1.14 (1.01) |
| Correct-estimators (N=8) | .42 (.26) | 1.30 (.28) | .06 (.30) | 1.33 (.77) |
| Under-estimators (N=10) | .48 (.32) | 1.70 (.45) | -.20 (.39) | .96 (1.01) |

*Note:* Parameter estimates derived from the Dual-Process Signal Detection model. Standard deviations are in parentheses.

### Response Times at Encoding

One possibility to account for the Dunning-Kruger Effect is that group-level differences could be due to how people encoded the information into memory (Craik & Lockhart, 1972; Addante et al., 2015); indeed, early accounts of the Dunning-Kruger Effect have posited that results can be due to competency of subjects that can be corrected by improving information acquisition (Kruger & Dunning, 1999). Similarly, subjects’ overconfidence at retrieval could have come from excess fluency at encoding providing feelings that the information was ‘easily learned’ (we thank an anonymous Reviewer for making this suggestion), leading them to rely upon intuitive perceptions of fluency and feelings of familiarity at retrieval that they incorrectly misattributed as better performance (Whittlesea & Leboe, 2000; Whittlesea & Leboe, 2003; Leynes & Zish, 2012; Leynes & Addante, 2016). To inform these possibilities, we analyzed mean encoding times as a general measure of information processing while participants encoded each item, using two-tailed between-group t-tests, and excluded one outlier in the group of over-estimators due to exceeding three standard deviations from the mean (Figure 4). Over-estimators (N = 37, *M* = 1289 ms, *SD* = 319, *SE* = 52) responded faster than under-estimators (N = 10, *M* = 1651 ms, *SD* = 655, *SE* = 207) during encoding (*t*(45) = −2.48, *p* = .016); Correct-estimators (N = 8, *M* = 1159 ms, *SD* = 331, *SE* = 117) also performed marginally faster than the under-estimators (*t*(16) = 1.92, *p* = .071). There were no reliable differences evident between RTs of over-estimators and correct-estimators (*t*(43) = 1.04, *p* = .303) (Figure 4). These findings appear to indicate that under-estimators may have performed better due to having spent (slightly) more time exposed to information tested later.

### Electrophysiological Results

The EEG data were analyzed in several systematic steps to probe possible differences between metacognitive judgements and cognitive strategies, and as noted in the Methods section, ERP analyses included only subjects who maintained a minimum number of valid ERP trials for both of the ERP conditions being compared, which resulted in somewhat smaller sample sizes from the original N = 61 and is noted in each reported result’s degrees of freedom. First, we assessed the data for general ERP differences that could be identified between the tasks of memory and metacognition judgments. To do this, we assessed the ERPs for decisions in all of the Dunning-Kruger judgments collapsed together compared to the ERPs for all item memory judgements collapsed together, to form a standard baseline for comparison since there had not been prior ERP work done in this kind of metacognitive task (Figure 5). This revealed that ERP activity for the metacognitive DKE decisions was significantly greater than those for memory judgements, starting from approximately 300 ms and continuing through 1000 ms at almost every electrode site. These effects were maximal at the central parietal site of Pz through 800 ms (300-500 ms: *t*(53) = 10.69, *p* < .001; 400-600 ms: *t*(54) = 15.19, *p* < .001; 600-900 ms: *t*(53) = 9.79, *p* < .001) and similarly reliable at several surrounding sites, upon which time the effects became evident as maximal at mid-frontal site Fz from 900-1200 ms (*t*(53) = 6.46, *p* < .001) with similar effects at surrounding sites, consistent with prior ERP findings of parietal and anterior P300a/b effects for novelty processing and oddball paradigms (Curran, 2004; Woodruff et al., 2006; Kishiyama et al., 2004; Knight, 1996; Knight & Scabini, 1998). This basic finding established a foundation that ERPs during the metacognitive judgments of the DKE were reliably distinct from memory-related activity, which we continued further investigation.

**Figure 5.**
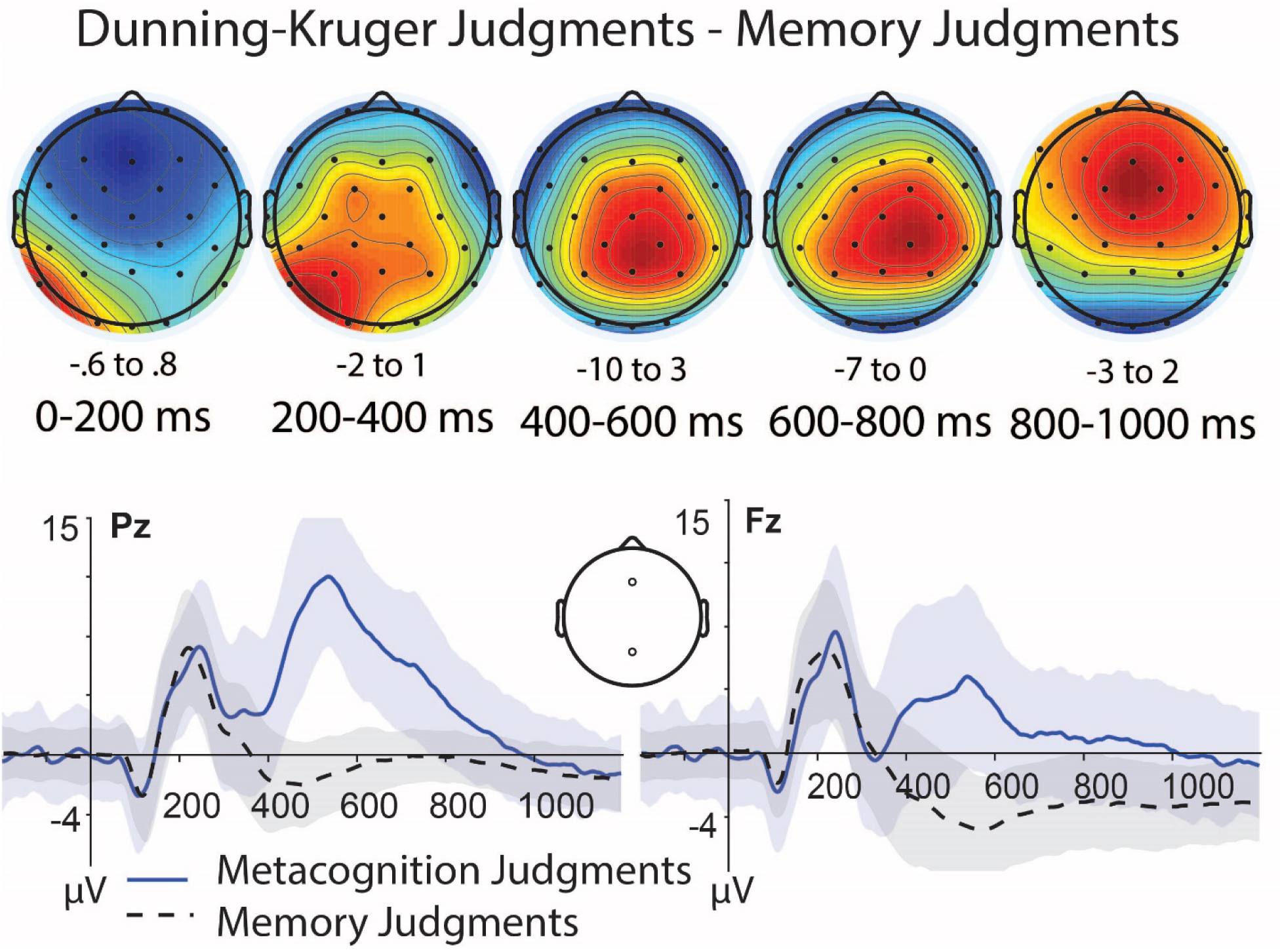
Comparison of ERPs for memory judgments vs metacognitive judgments estimating performance. a) Topographic difference maps of ERPs for all item memory judgments compared to all Dunning-Kruger judgments (DK judgments minus memory judgments). Each topographic map is range normalized according to their maximum and minimum values per latency. Warmer colors represent more positive-going voltage differences, with scales for each noted beneath each map. b) ERPs for memory and metacognition tasks at central parietal size Pz; x-axis is time in milliseconds, y-axis is µV. c) ERPs for memory and metacognition tasks at mid-frontal site Fz.

Are there differences in how DKE groups were making their memory judgments? We next investigated physiological differences in memory (‘old-new’ effects of hits - correct rejections) as a function of the different kinds of DKE groups (i.e., over-estimators, under-estimators). When looking early in time (400-600 ms) during the latencies characteristic of familiarity-based processing (FN400; Addante et al., 2012a,b; Rugg & Curran, 2007) a 2 x 2 repeated measures ANOVA with factors of Condition (ERP amplitudes for hits, correct rejections) and Group (over-estimators (N = 36), under-estimators (N = 10)) at mid-frontal site Fz revealed no significant effects of either factor or interaction. However, at the centro-parietal site Pz there was a significant main effect of Condition (*F*(1,46) = 5.63, p = .022) and a reliable condition x group interaction (*F*(1,44) = 5.0, p = .030). This was explored with a follow-up between-group t-test, Which found under-estimators had a significantly higher amplitude (*M* = 1.39, *SD* = 1.59) of old-new effects than over-estimators (*M* = 0.25, *SD* = 1.38) that occurred maximally at centro-parietal site Pz but was diffuse across several sites in the posterior scalp, *t*(44)= 2.24; *p* = .03. As this difference was evident in the parietal region instead of the expected left frontal region characteristic of the FN400, it may indicate an early activation of recollection activity but does not preclude other possible interpretations of its functional significance related to familiarity or implicit processing (Voss et al., 201; 2012; Voss & Paller, 2007; 2017; Addante, 2015).

Later in time, from 600-900 ms at left parietal site P3, a 2 x 2 repeated measures ANOVA with factors of Condition (ERP amplitudes for hits, correct rejections) and Group (over-estimators, under-estimators) from 600-900 ms at left parietal site P3 revealed a main effect of Condition (*F*(1,46) = 7.36, p = .009), and a Condition x Group interaction (*F*(1,44) = 9.91, p = .007). This finding was qualified by a between-group t-test, which revealed that the under-estimator group (*M* = 1.96, *SD* = 1.35) had significantly larger LPC effects (hit – correct rejection amplitudes) than the over-estimator group (*M* = 0.30, *SD* = 1.72), *t*(44)= 2.81; *p* = .01 (Figure 6). This finding was similar across adjacent electrodes, and suggests that the under-estimator group, which consists of the highest performing individuals, relied on using more recollection than the over-estimator group did in making their memory judgments. Accordingly, since the illusory over-estimators constituted the lowest performing individuals, it is possible that one reason why they performed lower was because of a relative reduction in their reliance upon recollection.

**Figure 6.**
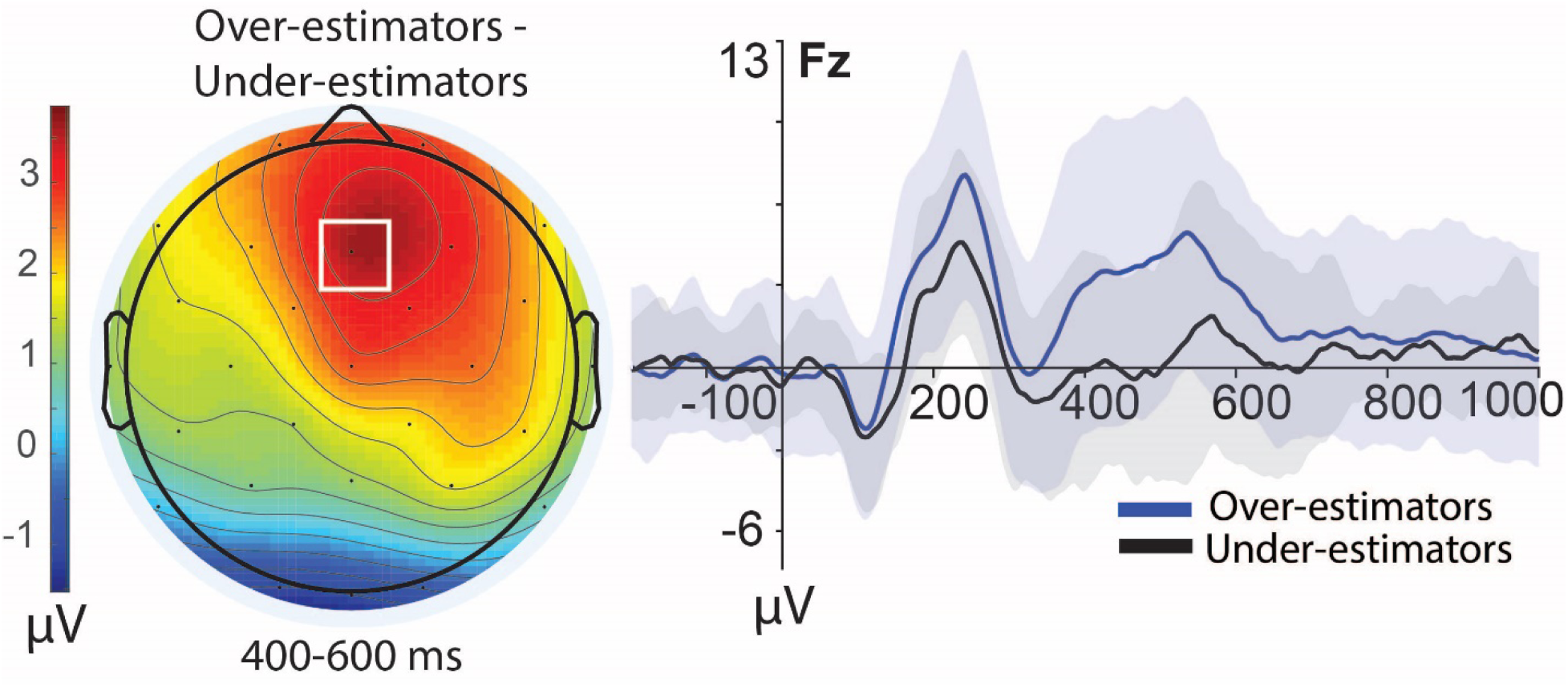
ERPs of Metacognitive Responses by Dunning-Kruger Groups. Topographic maps show ERPs of collapsed Dunning-Kruger responses (Dunning-Kruger judgments 1, 2, 3, 4, and 5 combined) for Over-Estimators compared to Under-Estimators from 400-600 ms. Topographic map is range normalized to maximum and minimum values, warmer colors represent more positive-going voltage differences. Right: ERPs for Dunning-Kruger judgments of over-estimators and under-estimators at mid-frontal site of Fz.

How do metacognitive judgments differ among good and bad, over- and under-estimators? To investigate this core question, we analyzed group-level differences in ERPs between the over- and under-estimators by DKE judgment (all responses collapsed together). There were significant differences in ERP amplitude between the under-estimators and over-estimators at mid-frontal electrode Fz from 400-600 ms (*M_Over-Estimators_* = 4.16, *SD* = 5.09; *M_Under-Estimators_* = 0.55, *SD* = 4.40; *t*(44) = −2.04, *p* = .048) and adjacent frontal sites, such that ERPs for over-estimators were far more positive than that of the under-estimators (Figure 7). One suggestion from these results is that the frontal effect at 400-600 ms may be characteristic of the FN400 ERP effect related to familiarity-based processing, in that over-estimators may be relying on the less-specific memory process of familiarity or intuitions of increased fluency to guide making their metacognitive judgments, instead of relying upon the more distinct recollection-related processes (Yonelinas et al., 2010) that evidently appears to be supporting the people who were under-estimating their performance relative to the group (Figure 7).

**Figure 7.**
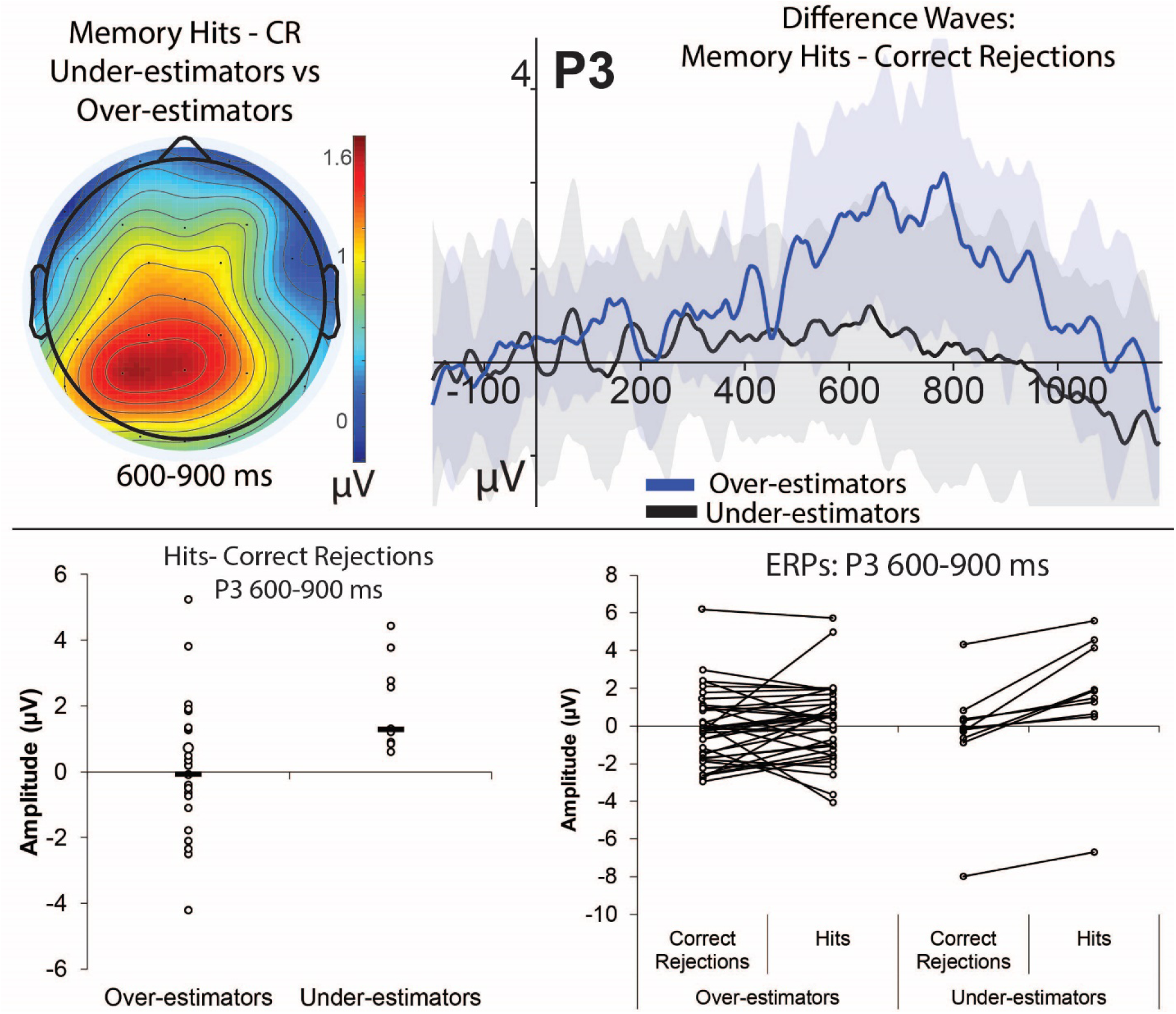
Difference waves of recognition memory ERP effects for Dunning-Kruger Groups. Top left: Topographic maps show group-difference waves for memory effects (hits minus correct rejections) for Dunning-Kruger groups of Over- and Under-Estimators at left parietal electrode P3, map is range normalized to maximum and minimum microvolts. Top right: ERPs of difference waves in memory effects (hits minus correction rejections) for each group (over- and under-estimators) at P3 from 600-900 ms; y-axis of zero represents no differences between memory conditions’ ERPs, and shaded areas depict standard error of the mean for each group. Warmer colors represent more positive-going voltage differences. Bottom left: individual raw amplitudes of the difference wave from 600-900ms, categorized by group. Bottom right: individual amplitudes of hits and correct rejections at P3 from 600-900ms for each subject, categorized by group.

### Brain-behavior Relationships Between Memory and Metacognition

The preceding results prompted the question of whether there is a systematic relationship evident between memory and metacognition effects at the individual subject level. Across subjects in groups of both over-estimators and under-estimators, the magnitude of the LPC effect that occurred between 600-900 ms at P3 (hits-CR) was found to be significantly correlated with the proportion of hits made by subjects (r = .318, *p* = .031, N = 46), and also negatively correlated with response speeds for times when high confidence hits went on to receive correct source memory responses (r = −.305, *p* = .039, N = 46) (Figure 8). For the under-estimator group, magnitude of the LPC effect was found to be correlated in the negative direction with the average Dunning-Kruger response given in-test by each subject (r = −.798, *p* = .006, N = 10) but did not exhibit any relationship for the over-estimators (r = −.014, *p* > .10, N = 36) (Figure 8), indicating that for the higher-performing under-estimators, LPC effects would reliably predict what the ensuing average Dunning-Kruger estimate of that subject would be for their estimated task performance, such that the larger LPC magnitudes predicted relatively lower performance estimates.

**Figure 8.**
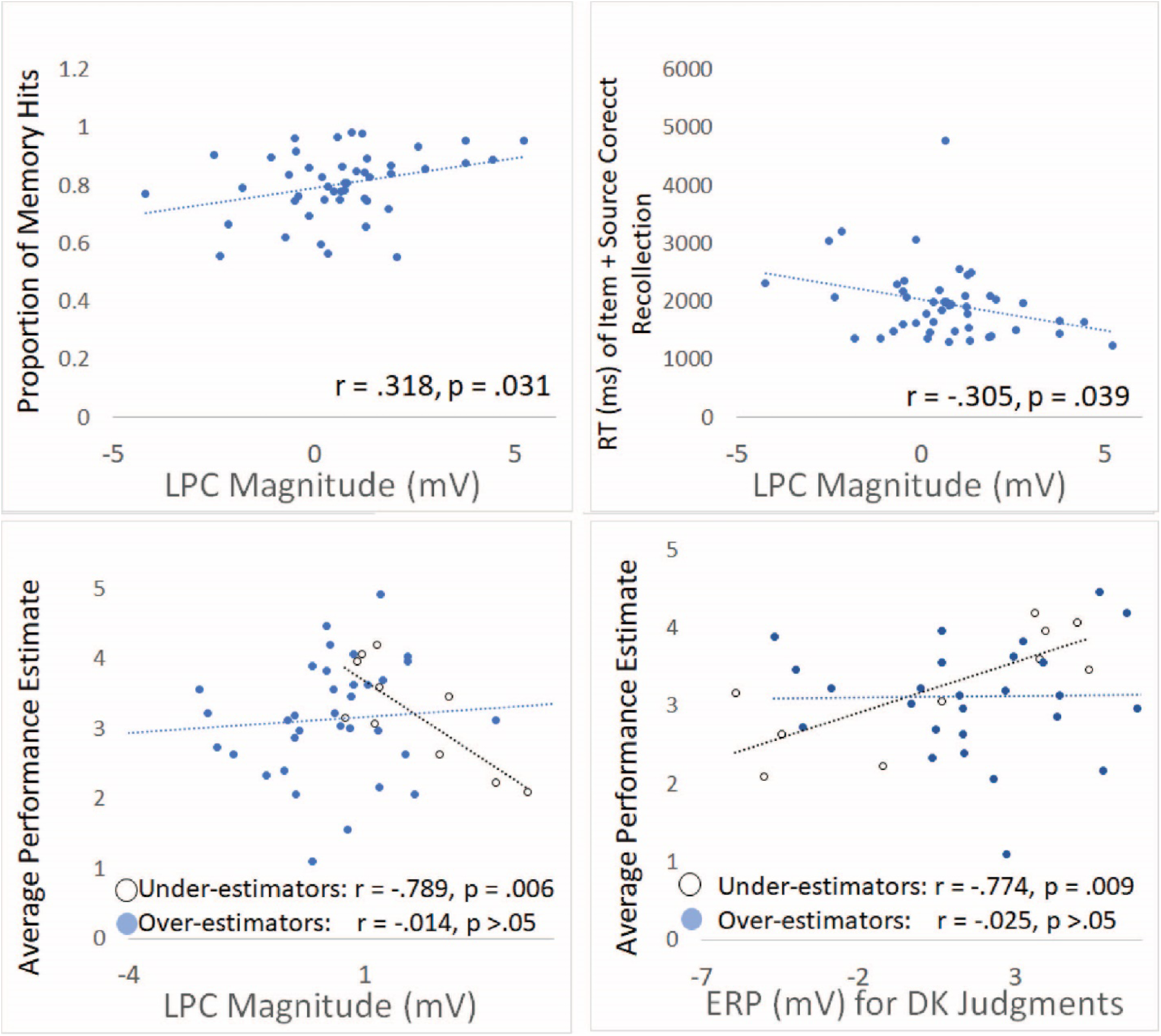
Relationships of Behavioral and Brain Measures for Memory and Metacognition. X-axis represents the magnitude of the LPC effect for both under-estimator and over-estimator groups combined (LPC measured as ERPs for hits minus correct rejections at left parietal site P3 from 600 – 900ms during item recognition memory test (top left, top right, bottom left panels). Bottom right panel x-axis represents the amplitude of mid-frontal ERPs for metacognitive judgments from 400-600ms during the in-test Dunning-Kruger performance estimate task, separated by group. Y-axis represents the proportion of successful item memory hits (judgments of 4 or 5 for ‘old’ status items during memory retrieval task) (top left); reaction times in milliseconds to recollection-related trials in which subjects got both an high item confidence hit and source memory judgment correct (top right); average in-test performance estimate given by subjects during the metacognitive DK judgments.

Overall, these findings converge to reveal that the larger LPC magnitudes were related to higher proportions of hits on the memory test and with faster response times for recollection-related items of high confidence hits with correct source memory (Addante et al., 2011; Addante et al., 2012a; Roberts et al., 2018). Hence, the LPC was related to recollection, and the more recollection signal (LPC) a subject had predicted the more likely they were to under-estimate their memory performance via their average DK responses in the metacognitive judgments (Figure 8). Recollection thus apparently led people to exhibit more humble metacognitive self-awareness. What is special about the under-estimators during their decisions to avoid the pitfalls of illusory superiority?

One line of evidence we found was that in the under-estimator group, the magnitude of the ERPs for metacognitive judgments from 400-600 ms at mid-frontal site of Fz exhibited a significant positive correlation with the average in-test Dunning-Kruger response given by subjects (r = .774, *p* = .009, N = 10), though this relationship was not evident for the over-estimators (r = .025, *p* = .886, N = 36) who exhibited the larger FN400-like effects (Figure 7, Figure 8). This suggested that the relative lack of familiarity-based processing in the under-estimators appears to be governing them towards reporting a lesser estimation of their performance in the task. That is, without the ambiguity of a familiarity signal it may leave recollection better suited to support these discriminating decisions. Overall, these findings indicate that we can reliably predict how people will perform in either accurately- or inaccurately-self-estimating their abilities by knowing the extent of their recollection-related brain activity occurring beforehand on a memory task (MacLeod & Donaldson, 2017; Addante et al., 2011).

## Discussion

The current study assessed multiple measures of Dunning-Kruger estimates interspersed throughout an on-going episodic memory test while EEG was recorded. The results from behavioral measures first revealed that the memory paradigm was successful at eliciting the DKE. Participants were separated into performance quartiles and their actual percentile ranking in the group was plotted alongside their estimated percentile ranking (Figure 2). The lowest performing participants in the bottom quartile were found to have substantially overestimated how highly they ranked in their groups while the highest performing participants moderately underestimated their actual ranking. This basic finding was important to identify as a starting point in a novel paradigm for studying the DKE in episodic memory, and its establishment permitted us to continue to explore the data in more specified ways for both behavioral and electrophysiological domains.

### Behavioral Findings

The current study’s paradigm permitted meaningful collection of reaction times for multiple Dunning-Kruger judgments that could be analyzed at a group level, which prior studies of the DKE have not been able to investigate due to their one-time measures of metacognitive performance estimates at the completion of a study. We found over-estimators were discernably faster than under-estimators in judging themselves to be in the top percentile, but they were slower to judge themselves as being in the bottom percentile; accordingly, under-estimators were relatively slower to report being in the best performance and quicker to claim they were doing poorly. There are several theoretical accounts that could be used to view these results, including elements of cognition and the traditional Dunning-Kruger account of double ignorance (1999).

The first account uses ‘cognition for prototypes’ to explain the reaction time patterns. Kruger and Dunning’s (1999) original results suggest that over-estimators do not understand that they are performing poorly and so they believe they are performing well and placing well within their participant group. This could lead to them having a very positive perception about their ability to perform well on certain types of tasks. Research on prototypes has shown that answers to questions that are very obviously true (closest to one’s prototype) are answered faster (for example, the question, “Is a robin a bird?” will elicit a faster “yes” response than the questions “Is an ostrich a bird?” even though both are true) (Rosch, 1974; Collins & Quillian, 1969). Therefore, if a person’s perception of their self (or prototype of themselves) includes that they perform well on tasks, they will be more likely to give a fast response when rating themselves well as opposed to rating themselves poorly. On the other hand, if they believe they are performing poorly, this perception would oppose the prototype that they have formed, causing them to react slower to rating their performance negatively. The same may be true if under-estimators have formed a perception about themselves that they are only average or even below average: it would then be logical that they would be slower to rate themselves as being best and quicker to believe they are performing poorly. Future research extending this paradigm with inclusion of ‘prototype’ measures of individual subjects would be able to test this hypothesis.

An alternative account of the reaction time results comes from using Kruger and Dunning’s (1999) model of double ignorance of low performers (i.e. 1. They do not know the answer, 2. They do not know they are ignorant of the answer) together with the inability of high performers to estimate their place among their peers due to not realizing the weaknesses of their peer group. By this account, over-estimators would be fast to report that they are doing well because they believe they are actually doing well, while they are slow to report that they are performing poorly because they do not believe they usually perform poorly or do not want to admit to themselves that they are performing poorly because that would be inconsistent with their metacognitive world-view.

Accordingly, the better-performing under-estimators would be competent enough to take pause in responding that they are doing well because they know the ways in which they might have failed (due to ‘competence’), and likewise also would be guided by a humble competence (knowing also what they may not know as well as they could know) to be quicker in believing they were performing poorly. The correlation analysis of RTs (Figure 8) suggest that these judgments may be based upon recollection processing evident from the electrophysiological effects.

### Neurophysiological findings

We began exploring the neurophysiology of the DKE by examining brain activity for general differences in processing among the memory and metacognition tasks. That is, we assessed the generic extent to which these two judgment types could be established for reflecting different kinds of processing. ERPs between memory trials and self-estimate trials were found to differ reliably beginning from approximately 300 ms into the epoch and continuing throughout the epoch to 1200 ms at almost every electrode site but being maximal first at posterior parietal sites and then later at mid-frontal regions (Figure 5). This pattern of ERPs is consistent with established properties of P300 ERP effects, or P3a and P3b effects, that are known to have the same distributions of topography and latency of across early/late and posterior/anterior regions, respectively, and which have been well-established as being associated with novelty processing or oddball tasks (Dien, Spencer, & Donchin, 2003; Otten & Donchin, 2009; Simons, et al., 2001). This is consistent with the current paradigm in that the DKE judgments were uncommon trials that appeared among the common memory trials in the test, and would have been salient stimuli for eliciting an orienting effect of attention as a novelty item (Kishiyama, Yonelinas, & Lazzara, 2006; Knight, 1996; Knight & Donatella, 1998).

We next explored whether the differential metacognitive judgments were associated with differential ERP patterns. When brain activity of all Dunning-Kruger responses was investigated together, over-estimators (those who performed worst) were found to have a higher mean ERP amplitude than under-estimators at frontal electrode sites during 400-600 ms (Figure 6, 8), consistent with the FN400 effect of familiarity-based processing (Addante et al., 2012a, b; for review see Rugg & Curran, 2007). These early ERP effects suggest that the errors of illusory superiority may be caused by an over-reliance on a generic sense of familiarity similar to as has been found in research on the False Fame Effect (Jacoby et al.,1989a, 1989b, 2004), as opposed to the more specific recollecting of the clear details from their past encounters which would instead provide the contextual cues to guide proper placement of one’s perceptual judgments. Under-estimators (those who performed best), on the other hand, exhibited a larger LPC than over-estimators did from 600-900 ms during memory judgments (Figures 7, 8), indicating that these under-estimators may be making the decisions by reliance upon the clearer details of recollected information, as opposed to the fuzzy sense of familiarity that can come with less accuracy (Yonelinas et al., 2004; 2010).

### A Memory-based Framework for Metacognitive Judgments

In the introduction we postulated that that a memory-based framework could account for the illusory errors seen in the DKE, whereby these errors (and successes) can be guided at least in part based upon differences in the cognitive processes of memory familiarity and recollection. In such a model, familiarity would be seen as providing the foundational cognitive processing associated with a heuristic used by people unsure about the details of their past performance on the task and thus guiding them to erroneously over-estimate how well they think they did (over-guessing based upon it feeling familiar but lacking details; Whittlesea, 2002; Whittlesea & Leboe, 2000; Whittlesea & Williams, 2000; Whittlesea, 1993; Whittlesea, Jacoby, & Girard, 1990; Whittlesea & Leboe, 2003; Voss & Paller, 2010). On the other hand, recollection would be seen as the cognitive process supporting people’s abilities to correctly retrieve their memories of the past experiences with richness, detail, and contextual information bound together with the item of the episode (Diana et al., 2008; Eichenbaum et al., 2007). Thus, having the cognitive process of recollection available would guide people to make self-assessments of performance that are more conservatively constrained by the details of the facts of that prior performance, thereby avoiding the same risk of incorrectly assuming an over-performance based merely on it seeming familiar acontextually. Taken together with the behavioral findings in RTs, it appears that over-estimators were ‘quick to brag’, whereas the high-performers were slow to judge themselves as being best- and their caution was associated with better scores.

Moreover, ERP data suggested that recollecting the past with clear context and details may be an important part to helping keep us humble, whereas relying upon mere feelings of familiarity (without being sure) may lead us to over-estimating ourselves.

What may instead delineate the matter is not confidence, per se, (since patients can be highly confident of familiar responses that lack both recollection and accuracy (Addante et al., 2012b, 2012a), but rather may be found in the construct of recollecting the fuller combination of items-bound-in-context (Diana et al., 2008; Eichenbaum et al., 2007; Addante et al., 2012a), including specific details, and contraindicating information (i.e. ‘non-criterial recollection’, Gallo, Weiss, & Schacter, 2004; Parks, 2007; Yonelinas & Jacoby, 1996) from which to place one’s judgment^2^.

Research on familiarity has identified a contribution of guessing, or fluency, to familiarity judgments that is included in its decision heuristic (Voss et al., 2010; Whittlesea, 2002; Whittlesea & Leboe, 2000; Whittlesea & Williams, 2000; Whittlesea, 1993; Whittlesea, Jacoby, & Girard, 1990; Whittlesea & Leboe, 2003) and that may evidently be leading people here to jump to the wrong conclusions about themselves relative to others, similar to as has been found in the False Fame Effect (Jacoby et al., 1989a, b, 2004). This concept of familiarity being involved in illusory superiority judgments builds from the ‘reach-around-knowledge’ account (Dunning, 2011), and the data here provide a more concrete substantiation of this within the framework of cognitive processes of episodic memory.

This interpretation is, to a certain extent, consistent with prior accounts of differences in people’s task competency (Kruger & Dunning 1999; Schlösser et al., 2013; Adams & Adams, 1960; Ehrlinger & Dunning, 2003; Oskamp, 1965; Pennycook et al., 2017; Ryvkin et al., 2012a; C. Sanchez & Dunning, 2017) if differences in illusory superiority judgments are understood as being due to differences in how people encoded the initial mnemonic information either successfully or unsuccessfully into memory. That is, those that did not encode information well (i.e. poor attention, motivation, or distraction; Craik, Luo, & Sakuta, 2010; Craik, Eftekhari, & Binns, 2018; Craik, Naveh-Benjamin, Ishaik, & Anderson, 2000; Fernandes, Moscovitch, Ziegler, & Grady, 2005; Galli, Gebert, & Otten, 2013; Addante et al., 2015; Middlebrooks, Kerr, & Castel, 2017; Weeks & Hasher, 2017) would not be likely to recollect that information later nor then be able to accurately calibrate how well they are actually performing if having to guess with heuristics of general familiarity and fluency (Whittlesea, 2002; Whittlesea & Leboe, 2000; Whittlesea & Williams, 2000; Whittlesea, 1993; Whittlesea, Jacoby, & Girard, 1990; Whittlesea & Leboe, 2003).

Accordingly, analysis of reaction times during encoding revealed that over-estimators responded faster than under-estimators while learning information, thereby having less time to encode and consolidate the items into memory (Figure 4), which is consistent with a large body of research on fluency in memory (Leynes & Zish, 2012; Bader & Mecklinger, 2017; Bruett & Leynes, 2015; Cermak, et al., 1992; Doss, Bluestone, & Gallo, 2016; Kurilla & Westerman, 2008; Leynes & Addante, 2016; Li, et al., 2015; Nie, et al., 2019; Thapar & Westerman, 2009; Volz, Schooler, & von Cramon, 2010; Westerman, 2008; Whittlesea & Leboe, 2000; Whittlesea & Leboe, 2003; Alter & Oppenheimer, 2009; Castel, McCabe, & Roediger, 2007; Hertzog, et al., 2003; Serra & Dunlosky, 2005). Thus, subjects could have responded more quickly to items at encoding by virtue of their seeming more fluent. While these findings appear to indicate that overconfidence may have stemmed from enhanced fluency at encoding leading one to believe the information was more easily learned, it is challenged by the finding of their having the same response time as the correct-estimators did. Alternatively, it appears that under-estimators may have performed better due to having spent (slightly) more time learning information better, again supporting their later competence. Future work will benefit from empirical manipulations of fluency (e.g.: Leynes & Zish, 2012; Bader & Mecklinger, 2017; Bruett & Leynes, 2015; Leynes & Addante, 2016) and physiological measures during the encoding.

### Alternative Interpretations

Studies of decision making have provided ERP evidence that P300 effect timing and slope is each associated with evidence accumulation in decision making tasks (O’Connell et al., 2012; Twomey et al., 2015; Boldt, et al., 2019). One possibility for the current results of group differences in ERPs during the performance estimates is that they may reflect differential decision making and evidence accrual among subjects (for a similar model, see Urai & Pfeffer, 2014) (we thank an anonymous Reviewer for pointing this out). By this account, over-estimators may have relied upon insufficient evidence accrual to make their hasty inaccurate decisions (consistent with the features of a familiarity-based signal detection process; Yonelinas et al., 2010; 2002), whereas the under-estimators may have been slower to believe they were doing best because of evidence accrual occurring more slowly for a slower-growing integration signal (Summerfield & Tickle, 2015; Twomey et al., 2015) (consistent with a threshold model of recollection; Yonelinas et al., 2010; 2002; Parks & Yonelinas, 2009; Yonelinas & Parks, 2007). The correlation results we found were consistent with this, in that larger P3 signal magnitudes for Dunning-Kruger decisions predicted higher performance estimates in the under-estimators, as they presumably had more accrued more evidence to support those judgments (Twomey et al., 2015; O’Connell et al., 2012; Boldt, 2019).

However, there are also a few lines of evidence weighing against this, which suggest that the results may not reflect core differences among groups in decision-making processes, attention to the task, or use of different strategies during the task. First, there were no differences across groups in their Dunning-Kruger response distributions, nor in their overall reaction times to the Dunning-Kruger decision task, which would be predicted by such accounts. Second, there were no differences across groups in quantification of their use of any decision-criterion shifts (C), nor sensitivity to response bias (B) (Table 6). Hence, while it is always possible there could be some other decision-making factor or differences that are driving the observed effects, none of the four direct measures of such indications revealed any evidence for it.

A extended possibility to that is that higher performing people might have gravitated to live in networks of other higher performing people that make them feel less outstanding (or at least, constrain their relative calibrations or criterions), and the lesser performing people might has socially gravitated to live in similar networks of lower performing people that permits them to presume having higher relative abilities. This possibility is speculative for now and lacks available data herein, and though intuitive it also runs counter to the current findings’ caution of relying upon intuition for making estimates of conclusions. While the current experiment is not equipped to fully explore these factors nor the roles of decision-making processes further, these interpretations present fruitful paths for future research to take these next steps of exploration.

### Implications

This current experiment provides several novel contributions to the understanding of the DKE that also leaves much room for future explorations to build upon. First, to our knowledge, this is the only Dunning-Kruger experiment in which self-estimates on a task relative to a peer group were recorded repeatedly throughout the task, and which include physiological measures. Self-estimates in prior studies are usually only acquired once: at the end of the task (Adams & Adams, 1960; Burson, Larrick, & Klayman, 2006; Ehrlinger & Dunning, 2003; Kruger & Dunning, 1999; Oskamp, 1965; Pennycook, et al., 2017; Ryvkin, Krajč, & Ortmann, 2012a; Sanchez & Dunning, 2017) (although see Simons (2013) for instance of when there was a variation of the task using repeated estimates before the task itself). This innovation to the classic Dunning-Kruger paradigm was critical to collecting both the reaction time measures and neurophysiological measures of ERPs during the metacognitive self-estimates, and offers future researchers a pathway forward to continuing exploring this phenomenon.

Our finding that under-estimators had a larger LPC than over-estimators did gives some insight into the inaccurate estimates of illusory superiority that occur in over-estimators (relative underperformers). Because the over-estimators had a smaller LPC (in fact, lacked any LPC evidence at all, Figure 7, 8), this suggests that they used less recollection during the episodic memory retrieval task. It is thereby reasonable to deduce that their memories for the episodic performance were diminished accordingly, leading to more metacognitive inaccuracies when trying to the recall episodic events related to their performance based purely upon the availability of familiarity. It is possible that this deficit may also be due to relatively more-impoverished encoding/learning in these subjects leading to relative differences in memory competency among the groups (Dunning, 2011), which can be tested in future studies.

We also found evidence of differences in brain activity between under-estimators and over-estimators when collapsing across all Dunning-Kruger metacognitive responses. Over-estimators had a larger ERP mean amplitude than under-estimators at mid-frontal electrode sites from 400-600 ms, which is the characteristic position and latency of the FN400 that has been synonymous with familiarity in many prior studies (Addante et al., 2012; Curran, 2000; Friedman, 2013; Gherman & Philiastides, 2015; Rugg et al., 1998; Rugg & Curran, 2007). In the framework of a memory-related interpretation of these results, one could argue that because we found an FN400 in this condition, over-estimators (under-performers) may have evidently used more familiarity than under-estimators did in making these metacognitive judgments, in lieu of the recollection that under-estimators (over-performers) were evidently relying upon instead. Evidently, each group was apparently arriving at fundamentally different metacognitive conclusions because they were relying upon, or being influenced by, fundamentally different neurocognitive processes of memory. This was mirrored in the behavioral data of reaction times, which revealed a crossover interaction pattern of responding: the over-estimators were quick to say they were best but slower to say they were worst, whereas the more ‘humble’ under-estimators were slower to say they performed best but faster to say they were worst-again suggesting fundamentally different cognitive processing between groups. This matched ERP findings that under-estimators relied upon recollection instead of familiarity, whereas those who assumed they were doing better than they were actually doing relied upon familiarty instead of recollection.

### Limitations and Considerations for Future Research

The current work maintains inherent limitations of all initial explorations: findings remain to be assessed for generalizability, tested for its boundary conditions, and independently investigated for replicability across other sample sizes and experimental variables. Some of our findings required relying upon relatively small sample sizes, and though the current paradigm has been previsouly found to be effective in seveal prior studies using even smaller sample sizes of neuropsychological patients (Addante et al., 2012; Addante, 2015), nevertheless, these findings should serve as preliminary discoveries to motivate future work exploring larger group sizes. It should be noted, though, that in attempting to characterize the effects of under-estimators, most people in the study exhibited over-estimating errors, so the majority of our relatievly large sized sample of N = 61 were defined as not being in the under-estimator group that avoided these errors (indeed, this more pervasive illusory error was the prime focus of the current study and maintained a relatively large ERP sample size). Specified analyses looked at conditions that were rarely responded to, which further reduced certain sample sizes. Nevertheless, in exploring these effects we maintained rigiorous controls of inclusion criteria for trials to gain effective signal to noise ratio (see Methods), as is attested to by the reliable effects observed in the current work that small sample sizes inherently create a stronger challenge to achieve (which we did).

Additionally, while the electrophysiological results are compelling in suggesting memory effects contributing to the Dunning-Kruger phenomena, we should be cautious to avoid an over-reliance upon reverse inference (Paller et al., 2012; Poldrack, 2011) since other cognitive processes also likely contribute to ERP effects of memory, too, such as implicit fluency and conceptual priming (Voss et al., 2014; Voss & Paller, 2010, 2012; Leynes & Zish; Leynes & Addante, 2016; though see comments in Addante et al., 2012a, 2012b; Mecklinger et al., 2012; Bridger et al., 2012; Bader et al., 2017). While future work would benefit from explorations in those directions, the current work is grounded in an extensive literature of established ERP findings (Rugg et a., 1998; Addante et al., 2012a; Addante et al., 2012b; for Review see Rugg & Curran 2007) and observed systematic relationships among behavioral and physiological correlates of the cognitve processes (Figure 8) (Stierset al., 2016; MacLeod & Donaldson, 2017). It also remains possible, though fully unexplored, that DKE for both behavior and physiology could co-vary on variables of personality, so futue work integrating combinations of electrophysiology, genetics, and personality inventories could prove to be fruitful. A final limitation to the current work is that the authors, too, may be inherently subject to the pervasiveness of the DKE and be over-estimating it’s value, misinterpreting results, or unaware of counterfactual evidence. We hope that the current research can serve in providing value for motivating future research investigating these findings in more depth, extend them, and test them against competing hypotheses.

### Summary and Conclusions

In conclusion, the current study adds to the literature by a series of small steps: first, it represents the first physiological measures of the DKE, as well as reaction time measures of the phenomenon. Second, the study represents an integrative new paradigm that was developed to permit measuring multiple recurring trials of Dunning-Kruger metacognitive judgments, which others can now use to extend our understanding further. Third, this paradigmatic innovation made possible the ability to capture the DKE in a complex episodic memory task which extends the body of work on the DKE to episodic memory tasks of item and source memory confidence measures/paradigms. Together, these innovations revealed convergent insight into why people differ in this phenomenon. People who made the illusory errors of over-estimating their performance did not have recollection-related processing supporting those memories and instead relied upon relatively less-accurate familiarity-based heuristics of intuition and fluency and were quicker to jump to those inaccurate conclusions. On the other hand, the more recollection signal (LPC) a subject had predicted the more likely they were to under-estimate their memory performance (Figure 9). Recollection thus appeared to lead people to a more cautious and constrained metacognitive comparison to others.

The current study thus represents a step forward in understanding one of the most pervasive observations of human behavior: metacognitive illusions from a tendency to both overestimate and underestimate our abilities relative to others. This pernicious psychological phenomenon has been observed by philosophers such as Socrates and Confucius (Socrates from Apology by Plato, 21d; Confucius, trans.

1938/500), is cautioned against by ancient texts including Judeo, Christian, Polynesian & Islamic traditions (Proverbs 12:15; 1 Corinthians, 3:18; Qur’an 31:18), noted by laureates (Shakespeare) & scientists (Charles Darwin) alike, and persists today throughout the modern age-including throughout university professors, provosts, deans, and peer-review (Cross, 1977; Huang, 2013) while extending to leaders occupying both the highest and lowest offices. The basic premise of the DKE is thus seemingly a fundamental force that shapes our socio-psychological universe in similar ways that gravity shapes the backdrop of our physical universe-persisting through time and affecting everyone at some level. It takes work with self-awareness to avoid the pitfalls of illusory superiority, and surely benefits from practice. We show here that one way to do that is to avoid relying on intuition, fluency, and familiarity to make quick judgments; instead, results encourage relying on recollection of details and slower responses to reduce errors of illusory superiority. More experimentation is needed, but the present work identifies some of the cognitive processes involved in the errors that can lead to the leadership and safety hazards of over- and under-confidence in one’s abilities. We hope that this research can serve to inspire new explorations endeavoring to discover the neural correlates of our psychological processes, towards a better understanding of ourselves and the truth of human behavior.

## Acknowledgements

Authors would like to thank Rose DeKock, Raechel Marino, Constance Greenwood, Celene Gonzalez, Yoselin Canizales, Kevin Benitez, Yesensia Casas, Maynori Hinton, and Roman Lopez for helpful assistance in collecting the EEG data; Rosalinda Valencia for assitance with analyzing the encoding reaction times; Dr. Joshua Koen for assitance with the ROC Toolbox used for calculating parameter estimates; Dr. Andrew Leynes for helpful discussions about effects; Drs. Donna Garcia and John Clapper for thoughtful feedback on an earlier draft, and are grateful to two anonymous Reviewers for insightful comments and recommendations. This research was supported by the following grants for which the authors acknowledge appreciation for the generous support: National Institute of Health Grant 1 L30 NS112849-01 to RJA from National Institute of Neurological Disorders & Stroke; The Mini-Grant Program from the CSUSB Office of Sponsored Research to RJA; the Faculty Assigned Time Grants from the CSUSB Office of Student Research to RJA; the Faculty Summer Research Fellowships from the CSUSB Office of Academic Research & the Deans Summer Research Fellowship from the College of Social & Behavioral Sciences to RJA; CSUSB Assigned Time for Exceptional Service to Students Award to RJA from Faculty Senate & Provost; the Faculty-Student Research Grants to RJA, LAS, & AM; the CSUSB ASI Student Travel Grants to AM and LAS; the CSUSB Student Success Initiative Innovative Scholars Fund to LAS and AM; and the CSUSB Student Success Initiative Culminating Project Awards to LAS and AM, and the Outstanding Master’s Thesis Award to AM from the CSUSB Office of Graduate Studies.

## Conflict of Interest Statement

Authors report no conflicts of interests.

## Data Accessibility Statement

Data is accessible upon request.

## Author Contributions

AM collected the data for a Master’s Thesis, analyzed data, and co-wrote the manuscript LAS programmed the study, assisted with behavioral data analysis RJA designed the study, supervised all parts, analyzed data, wrote the manuscript, handled the submission process, and revised the manuscript for resubmission.

## Abbreviations

DKE: Dunning-Kruger Effect
EEG: Electroencephalogram
ERP: Event-related Potential

## Footnotes

We used the post-test relative Dunning-Kruger estimate, so as to be consistent with the original approach used in Kruger & Dunning (1999), although we also conducted a paired t-test between the average of the in-test Dunning-Kruger responses (*M* = 3.14, *SD* = 0.81) for each person to the post-test relative Dunning-Kruger response (*M* = 3.16, *SD* = 0.78) and found that the two scores did not differ, *t*(55) = 1.30, *p* = .200.

Though note, that simply being ‘sure’ is not sufficient on its own, either; see Introduction review of the anti-correlation of memory and confidence, since success is not guaranteed with mere high confidence (Chua, Hannula, & Ranganath, 2012; Roediger & DeSoto, 2014; Koriat et al., 2008; Nelson & Narens, 1990; Wells, et al., 2002).

## Notes

### Competing Interest Statement

The authors have declared no competing interest.

### Summary of Updates

Text has been revised with additional analyses, clarifications, and with figures & interpretations updated accordingly.

